# Repeated Omicron exposures override ancestral SARS-CoV-2 immune imprinting

**DOI:** 10.1101/2023.05.01.538516

**Authors:** Ayijiang Yisimayi, Weiliang Song, Jing Wang, Fanchong Jian, Yuanling Yu, Xiaosu Chen, Yanli Xu, Sijie Yang, Xiao Niu, Tianhe Xiao, Jing Wang, Lijuan Zhao, Haiyan Sun, Ran An, Na Zhang, Yao Wang, Peng Wang, Lingling Yu, Zhe Lv, Qingqing Gu, Fei Shao, Ronghua Jin, Zhongyang Shen, Xiaoliang Sunney Xie, Youchun Wang, Yunlong Cao

## Abstract

The continuous emergence of highly immune evasive SARS-CoV-2 variants, like XBB.1.5^1,2^ and XBB.1.16^3,4^, highlights the need to update COVID-19 vaccine compositions. However, immune imprinting induced by wildtype (WT)-based vaccination would compromise the antibody response to Omicron-based boosters^5-9^. Vaccination strategies that can counter immune imprinting are critically needed. In this study, we investigated the degree and dynamics of immune imprinting in mouse models and human cohorts, especially focusing on the role of repeated Omicron stimulation. Our results show that in mice, the efficacy of single Omicron-boosting is heavily limited by immune imprinting, especially when using variants antigenically distinct from WT, like XBB, while the concerning situation could be largely mitigated by a second Omicron booster. Similarly, in humans, we found that repeated Omicron infections could also alleviate WT-vaccination-induced immune imprinting and generate high neutralizing titers against XBB.1.5 and XBB.1.16 in both plasma and nasal mucosa. By isolating 781 RBD-targeting mAbs from repeated Omicron infection cohorts, we revealed that double Omicron exposure alleviates immune imprinting by generating a large proportion of highly matured and potent Omicron-specific antibodies. Importantly, epitope characterization using deep mutational scanning (DMS) showed that these Omicron-specific antibodies target distinct RBD epitopes compared to WT-induced antibodies, and the bias towards non-neutralizing epitopes observed in single Omicron exposures due to imprinting was largely restored after repeated Omicron stimulation, together leading to a substantial neutralizing epitope shift. Based on the DMS profiles, we identified evolution hotspots of XBB.1.5 RBD and demonstrated the combinations of these mutations could further boost XBB.1.5’s immune-evasion capability while maintaining high ACE2 binding affinity. Our findings suggest the WT component should be abandoned when updating COVID-19 vaccine antigen compositions to XBB lineages, and those who haven’t been exposed to Omicron yet should receive two updated vaccine boosters.

## Main

SARS-CoV-2 continues to evolve, and new mutants constantly emerge under humoral immune pressure^10-14^. New variants, such as the XBB lineages, are capable of evading antibodies induced by vaccination or infection, resulting in repeated infections among populations^5,7,15,16^. Therefore, it is critical to develop updated vaccines that can elicit strong immune responses against the latest variants.

mRNA vaccine platforms can quickly adapt to new SARS-CoV-2 variants^17-20^. However, since the majority of the population was vaccinated with the ancestral SARS-CoV-2 strain, immune imprinting induced by WT vaccination presents a major challenge to the performance of updated boosters^21,22^. This is because boosting with a variant antigenically distinct from WT would majorly recall memory B cells induced by WT vaccination and masks the *de novo* generation of variant-specific B cells, which would hinder the generation of appropriate humoral immunity against new and emerging variants^6,7,9,23-27^.

It is crucial to explore vaccination strategies that can counter immune imprinting. In this paper, we investigated the dynamics of immune imprinting in both mouse models and human cohorts, with a particular focus on how repeated exposure to Omicron variants could alleviate immune imprinting.

### Alleviation of immune imprinting in mice

First, we investigated the effects of WT-vaccination-induced SARS-CoV-2 immune imprinting in mice. To accomplish this, two doses of 3 μg CoronaVac (an inactivated vaccine derived from the wildtype SARS-CoV-2) were used as primary immunization, and variant Spike proteins were used as boosters^28-30^. All SARS-CoV-2 Spike proteins contained six proline substitutions (S6P) and alanine substitutions in the furin cleavage site to stabilize in prefusion conformation^31^.

Mice that received a single booster of 10 μg Spike protein, including BA.1, BA.5, BQ.1.1, XBB, and SARS-CoV-1, showed decreased serum 50% neutralizing titers (NT50s) (VSV-based pseudovirus) against the D614G as the antigenic distance between the boosting variant and the wildtype increased, suggesting decreased cross-reactive B cell recall after the variant booster (Fig. 1a). Additionally, single dose boosted mice had significantly lower NT50 against the boosting variants compared to D614G (Fig. 1a). These results revealed substantial ancestral strain immune imprinting at the serum level, and are consistent with the observations in humans^6,7,23,24,32,33^, as well as previous findings of immune imprinting in influenza viruses^34,35^.

**Fig. 1.**
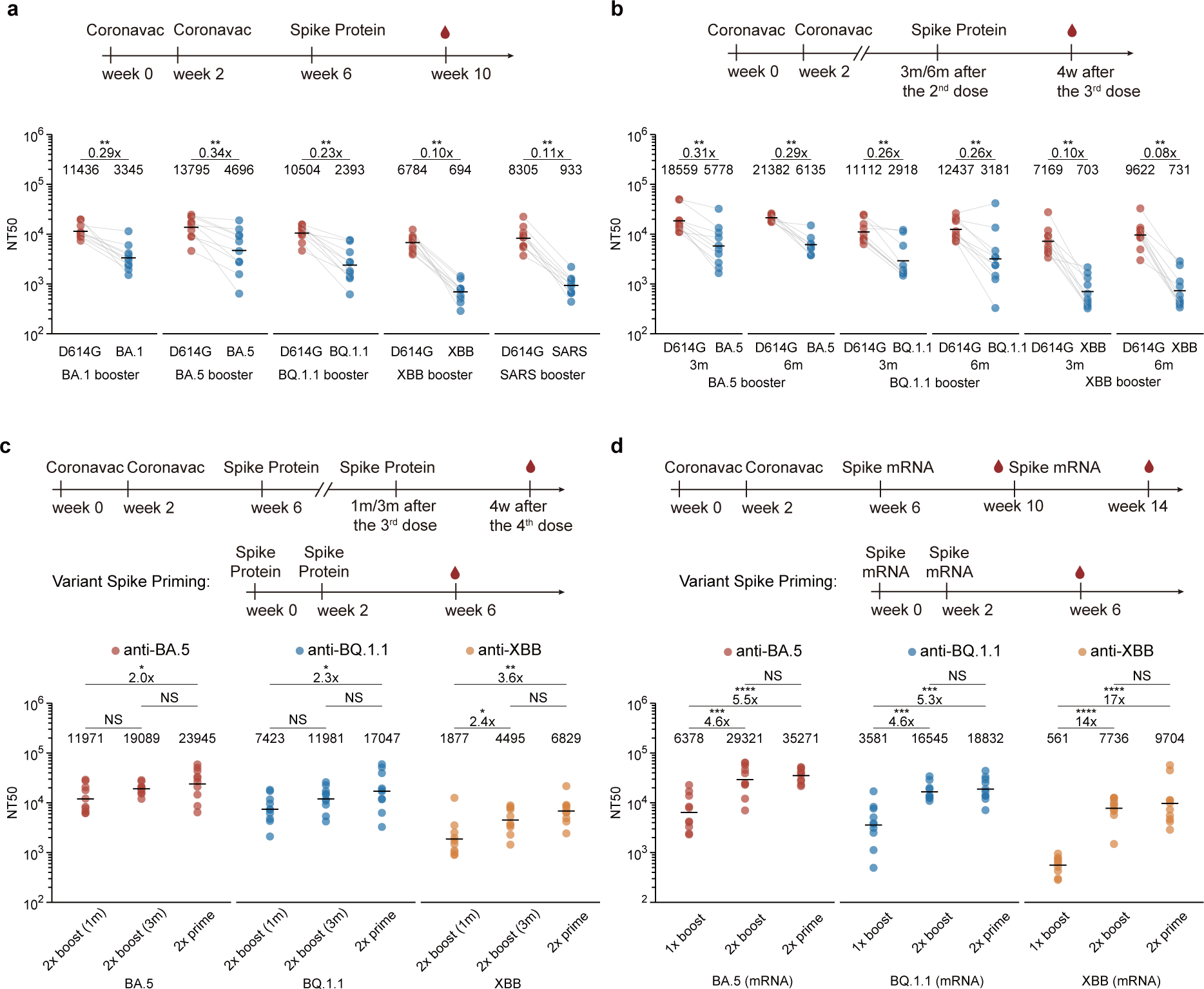
Humoral immune imprinting in mice. **a**, Neutralizing antibody response after priming with 2 doses of 3 μg CoronaVac followed by boosting with 10 μg SARS-CoV-1 Spike protein or SARS-CoV-2 variant Spike proteins in mice. **b**, Neutralizing antibody response after immunization with 2 doses of 3 μg CoronaVac followed by boosting with 10 μg SARS-CoV-2 variant Spike proteins with 3-month or 6-month time intervals in mice. The variants labled on x-axis of the graphs indicate the NT50 against that variant in (**a**, **b**). The variants marked at the bottom of the figure are the variants used for boosting in **(a, b)**. **c**, Neutralizing antibody response after priming with 2 doses of 3 μg CoronaVac followed by boosting twice with 10 μg SARS-CoV-2 variant Spike proteins with 1-month or 3-month intervals in mice. **d**, Neutralizing antibody response after priming with 2 doses of 3 μg CoronaVac followed by boosting twice with 1 μg SARS-CoV-2 variant Spike mRNAs. The variants marked at the bottom of the figure are the variants used for priming or boosting in (**c**, **d**). Red, blue, yellow circuls indicate the NT50s against BA.5, BQ.1.1, and XBB in (**c**, **d**). 10 mice were immunized and analyzed in each group (n= 10). Sera were collected four weeks after the last dose. Geometric mean titers (GMT) were labeled. For paired samples in **a**-**b**, statistical significance was determined using two-tailed Wilcoxon signed-rank tests. For independent samples in **c**-**d**, statistical significance were determined using two-tailed Wilcoxon rank sum tests. *p < 0.05, **p < 0.01, ***p < 0.001, **** p < 0.0001, and not significant (NS) p > 0.05. All neutralization assays were conducted in at least two independent experiments.

To investigate whether prolonging the interval between the primary WT immunization and the variant booster could alleviate immune imprinting, we further tested boosting mice 3-month and 6-month after CoronaVac priming (Fig. 1b). It was observed that 3-month and 6-month intervals between WT-priming and variant-boosting slightly increased overall NT50s, but the fold-change between NT50s against D614G and XBB remained high (Fig. 1b). Moreover, no significant NT50s difference among 1-month, 3month, and 6-month boosting groups was observed for BQ.1.1 and XBB boosting(Extended Data Fig. 1a-b). This suggests that longer intervals between the priming and Omicron-boosting, which would allow the maturation of WT-induced antibodies, may not be sufficient to alleviate immune imprinting.

The efficacy of the first Omicron booster is heavily limited by immune imprinting. It’s crucial to examine how a second Omicron booster performs^36^. We started by boosting CoronaVac-primed mice with two doses of the variant Spike protein over a 1-month or 3-month interval (Fig. 1c). Importantly, the second boosters resulted in increased NT50s against the corresponding variants (Extended Data Fig. 2a), as well as substantially reduced fold-changes between the D614G and variants (Extended Data Fig. 2b). However, the neutralizing titers induced by two boosters over one-month interval after two doses of CoronaVac priming were still lower than those induced by two doses of variant priming, clearly indicating the interference caused by immune imprinting (Fig. 1c). Notably, compared to 1-month boosting interval, a 3-month interval between Omicron boosters resulted in clear improvements in NT50s against all the corresponding boosting variants (Fig. 1c), and the fold-change between the NT50s against D614G and the boosting variants also decreased (Extended Data Fig. 2b). This indicates that the maturation of B cells induced by Omicron-boosting are highly beneficial for immune imprinting mitigation.

**Fig. 2.**
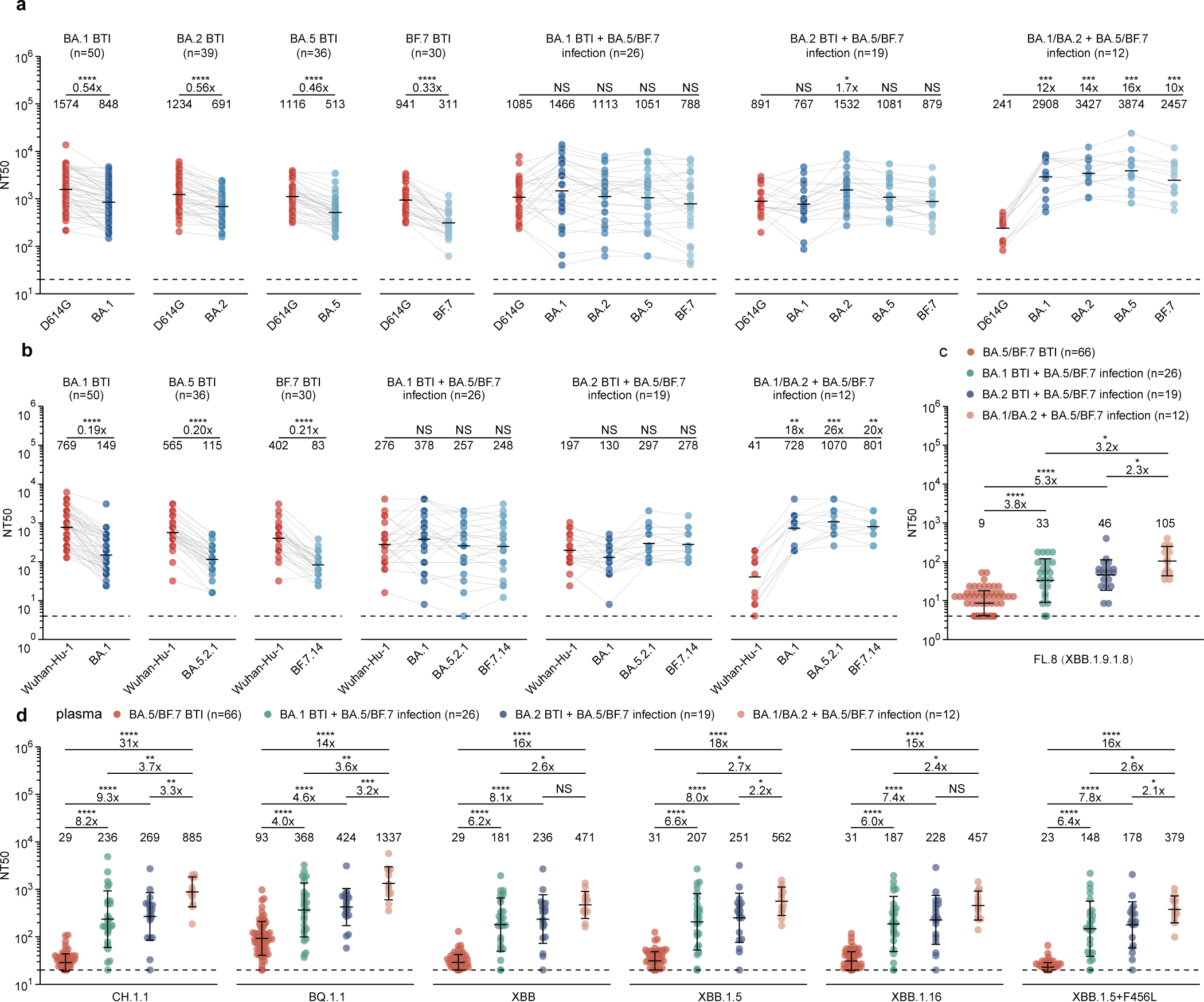
Humoral immune imprinting after repeated Omicron infections in humans. **a**, Examination of immune imprinting after Omicron breakthrough infections and repeated Omicron infections. Plasma antibody titers against pseudotyped D614G and variants were measured. **b**, Plasma antibody titers against authentic virus variant. For (**a**, **b**), fold changes between titers against variants and D614G were calculated and shown above the line. Statistical significance was determined using the Wilcoxon signed-rank test. **c**, Plasma antibody titers against authentic FL.8 (XBB.1.9.1.8) after repeated Omicron infections and BA.5 or BF.7 breakthrough infections. Fold changes between titers of different cohorts were calculated and shown above the line. Statistical significance was determined using the Wilcoxon rank sum tests. **d**, Plasma antibody breadth after one-time breakthrough infection and repeated Omicron infections. Plasma antibody titers against circulating pseudotyped variants were measured. Fold changes between titers of different cohorts were calculated and shown above the line. Statistical significance was determined using the Wilcoxon rank sum tests. BA.1, BA.2, BA.5, BF.7 BTI: post-vaccination Omicron breakthrough infection (BTI). BA.1, BA.2 BTI+ BA.5/BF.7 infection: post-vaccination Omicron breakthrough infection followed by BA.5/BF.7 reinfection. BA.1/BA.2+ BA.5/BF.7 infection: BA.1/BA.2 infection followed by BA.5/BF.7 reinfection with no vaccination history. Blood samples were collected 1-2 months after the last infection. Detailed information about the cohorts is in Supplementary Table 1. Geometric mean titers (GMT) are labeled in (**a**, **b**). Geometric mean ± SD are labeled in (**c**-**d**). Dashed lines indicate the limit of detection (LOD, NT50 = 20). *p < 0.05, **p < 0.01, ***p<0.001, ****p<0.0001, and not significant (NS) p > 0.05. All neutralization assays were conducted in at least two independent experiments.

Since mRNA vaccines encoding Spike have proved to be capable of quick adaptation to new variants, it is critical to test how updated mRNA variant boosters perform, especially when the higher immunogenicity of mRNA vaccine might help alleviate immune imprinting when served as Omicron boosters. Therefore, we tested 1 μg mRNA vaccines encoding BA.5, BQ.1.1, and XBB Spike as boosters in replacement of protein boosters (Fig. 1d). As expected, 1 μg mRNA vaccine demonstrated higher immunogenicity than the protein vaccine (Extended Data Fig. 2c-d, f). However, the performance of one-dose mRNA Omicron-boosters is still heavily interfered by immune imprinting despite higher immunogenicity, while two mRNA Omicron-boosters would significantly increase antibody titers and could achieve similar titers compare to the priming groups (Fig. 1d and Extended Data Fig. 2c-e). This suggests raising the immunogenicity of variant boosters could help to counter immune imprinting brought by WT vaccination.

Notably, among the Omicron variants tested, XBB boosting exhibited the lowest overall titers (Fig. 1c-d). Indeed, these variant vaccines, whether protein or mRNA, exhibit different levels of immunogenicity in mice, with XBB demonstrating the lowest (Extended Data Fig. 2f).

Together, our results observed in mice emphasize that the efficacy of the first Omicron boosters is severely limited by immune imprinting while a second booster is almost mandatory to alleviate immune imprinting and generate high antibody responses, especially for boosters encoding variants that exhibit long antigenic distance from WT, such as XBB.

### Mitigating immune imprinting in humans

To verify whether the findings obtained from mice also apply to humans, we recruited cohorts with repeated Omicron breakthrough infections (BTIs), including individuals with post-vaccination BA.1 or BA.2 BTI followed by BA.5/BF.7 reinfection (BTI+reinfection) and compared them to previously reported BA.1, BA.2, BA.5, BF.7 one-time BTI cohorts^7,32,37,38^. Importantly, we also included individuals who had no history of SARS-CoV-2 vaccination before repeated infection (vaccination-naïve reinfection) as controls. Detailed information about the cohorts can be found in Supplementary Table 1. We first tested neutralizing titers against exposed variants of these cohorts with pseudovirus and authentic virus neutralizing assays (Fig. 2a-b). Similar to mice immunization results, plasma neutralizing titers induced by one-time Omicron BTIs against the corresponding variant were significantly lower than those against D614G, consistent with our previous report^7^, and the fold changes between the NT50 against D614G and those against corresponding variants also increased as the antigenic distance increases (Fig. 2a-b). As expected, in the repeated Omicron infection group, with or without SARS-CoV-2 vaccination history, the neutralizing titers against Omicron variants significantly increased compared to one-time BTIs (Fig. 2a-b). More importantly, BA.1 or BA.2 BTI followed by BA.5/ BF.7 reinfections demonstrate comparable NT50 between exposed Omicron variants and D614G, indicating immune imprinting alleviation by the second Omicron exposure (Fig. 2a-b). However, the NT50s of vaccination-naïve reinfection group against Omicron variants were the highest among these cohorts (Fig. 2a-b), suggesting that repeated BTIs were still subjected to WT-vaccination-induced immune imprinting. Compared to one-time BTIs, repeated Omicron infection also led to an increase in the neutralizing titers against highly immune-evasive CH.1.1, BQ.1.1, XBB, FL.8 (XBB.1.9.1.8), XBB.1.5, XBB.1.16, and XBB.1.5+F456L (Fig. 2c-d and Extended Data Fig. 3a-c), indicating that repeated Omicron infections may broaden the breadth of antibody response. In addition, we found that the nasal swab samples from individuals with repeated Omicron infection exhibited higher neutralizing titers against Omicron variants than one-time breakthrough infection, suggesting strong nasal mucosal humoral immunity has been established after repeated infection (Extended Data Fig. 4).

**Fig. 3.**
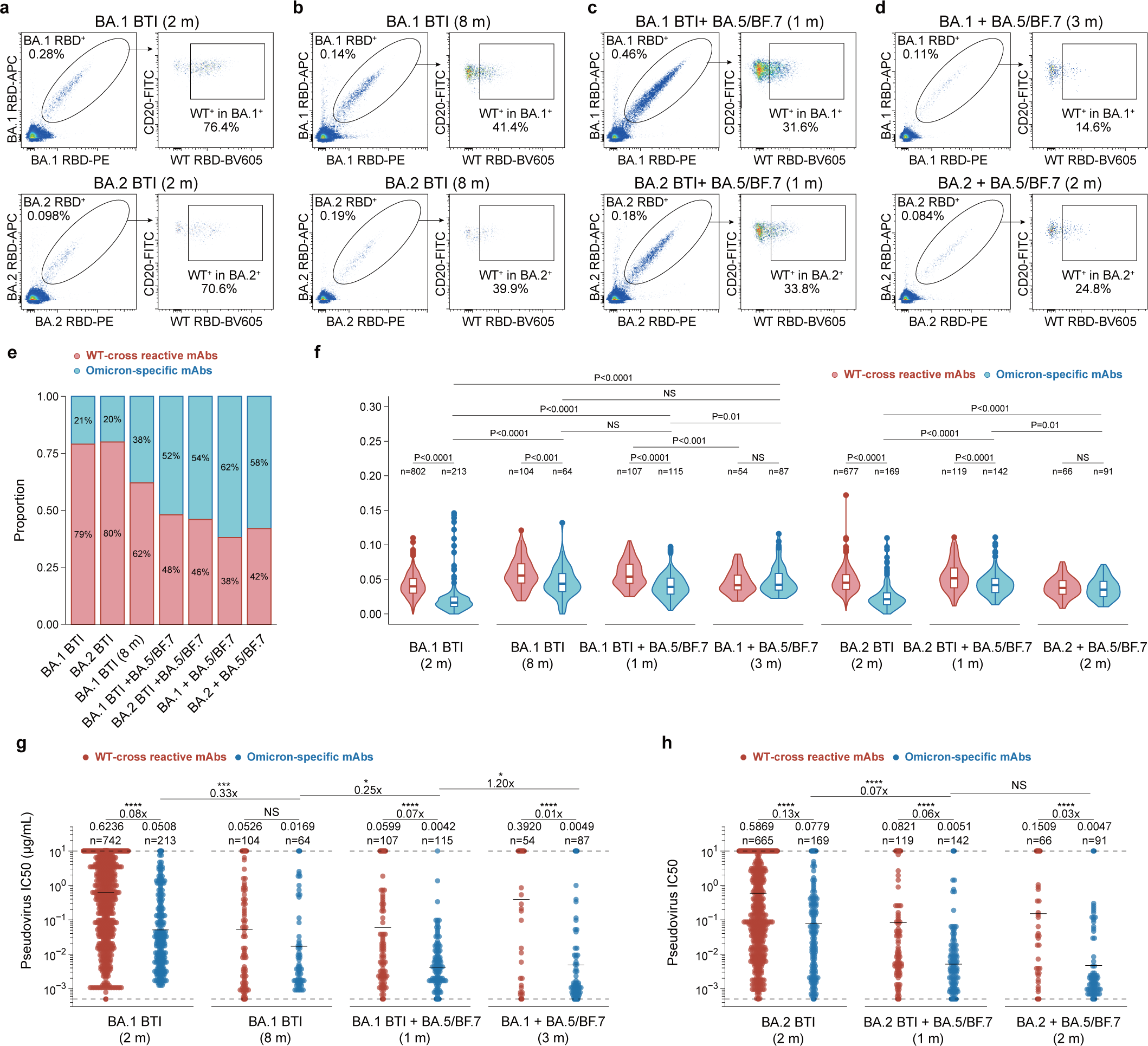
B cell immune imprinting after repeated Omicron infections. **a–d**, Flow cytometry analysis of pooled B cells from Omicron infection convalescent individuals. BA.1 (up) and BA.2 (down) RBD double-positive CD20+, IgM-, IgD-, CD27+ B cells were isolated for paired-single-cell V(D)J sequencing. Flow cytometry analyses were performed in cohorts of the following: (**a**) 2 months after BA.1 (up) or BA.2 (down) breakthrough infections, (**b**) 8 months after BA.1 (up) or BA.2 (down) breakthrough infections, (**c**) 1 month after BA.5/BF.7 reinfection after BA.1 (up) and BA.2 (down) breakthrough infections, **(d**) 2-3 months after BA.5/BF.7 reinfection after BA.1 (up) or BA.2 (down) infection without SARS-CoV-2 vaccination history. APC, allophycocyanine; FITC, fluorescein isothiocyanate; PE, phycoerythrin. BV605, Brilliant Violet 605. **e**, Proportions of WT-binding and non-WT-binding antibodies from Omicron breakthrough infection and repeated Omicron infection cohorts. Binding specificity was determined by ELISA. The antibodies were expressed in vitro using the sequence of the RBD-binding memory B cells from various cohorts. **f**, The heavy-chain variable domain somatic hypermutation rate of the mAbs from various cohorts. Statistical tests were determined using two-tailed Wilcoxon rank-sum tests. Boxes display the 25th percentile, median and 75th percentile, and whiskers indicate median ± 1.5 times the interquartile range. Violin plots show kernel density estimation curves of the distribution. The numbers and ratios of samples in each group are labeled above the violin plots. **g-h,** The BA.1(g) or BA.2(h) pseudovirus neutralizing ability(IC50) of the mAbs from various cohorts. Detection limit is denoted as dashed line, and geometric mean is denoted as black bar. Geometric mean, fold changes and the number of antibodies are labeled above the plots. Statistical tests were determined using two-tailed Wilcoxon rank-sum tests in (**f-h**). *p < 0.05, **p < 0.01, ***p<0.001, ****p<0.0001, and not significant (NS) p > 0.05.

**Fig. 4.**
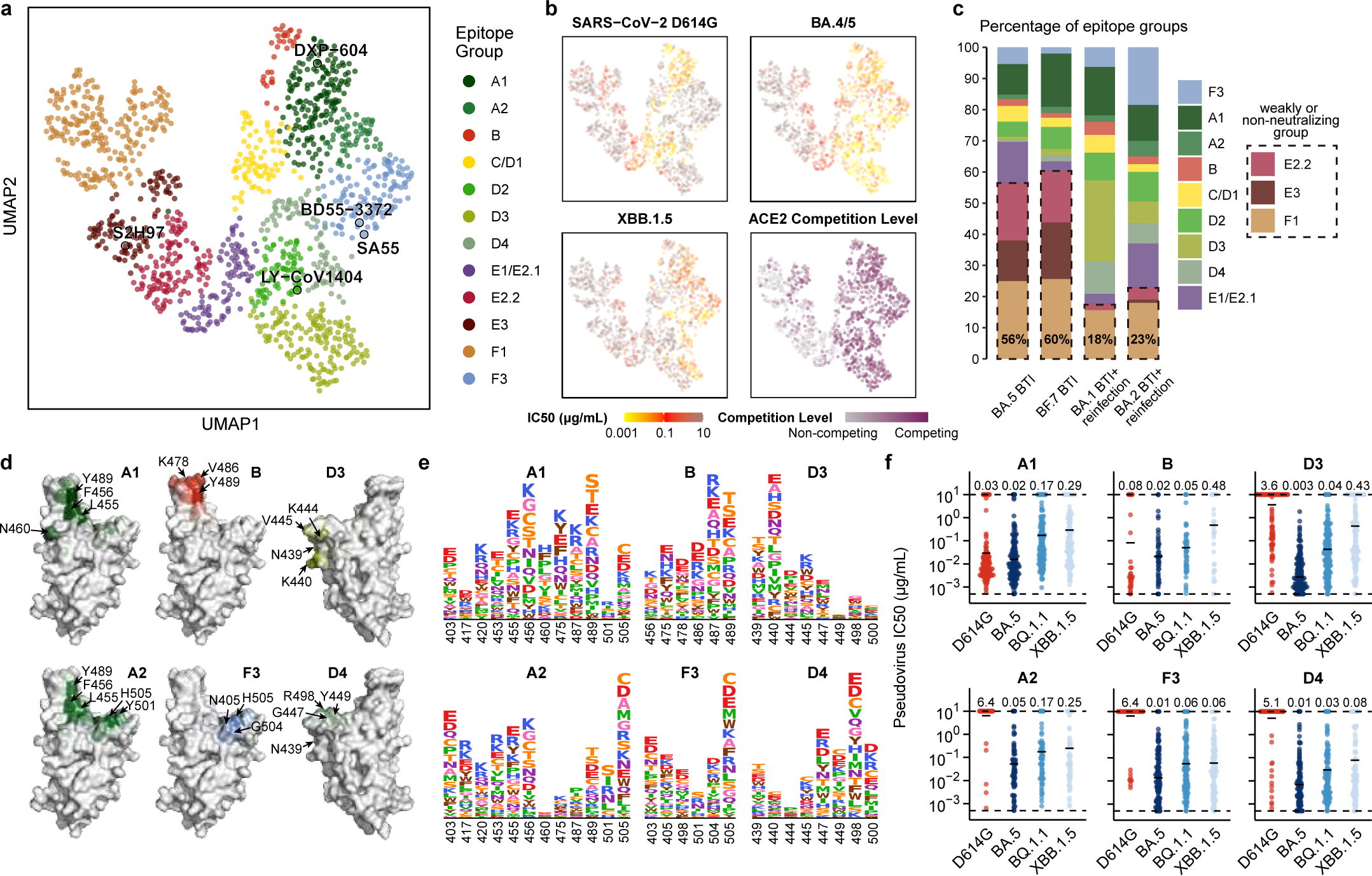
Epitope distribution and characterization of mAbs elicited by Omicron BTI and reinfection. **a**, UMAP embedding of epitope groups of monoclonal antibodies (mAbs) binding BA.5 RBD isolated from convalescent individuals who experienced BA.5/BF.7 BTI or reinfection (n=1350). **b**, Neutralization activities, denoted as IC50 values, against SARS-CoV-2 D614G (n = 1349), BA.4/5 (n = 1322), and XBB.1.5 (n = 1346) spike-pseudotyped vesicular stomatitis viruses (VSV), as well as ACE2 competition levels determined by ELISA (n = 1344), are projected onto the UMAP embedding space. **c**, Distribution of mAbs across epitope groups is shown for BA.5 Breakthrough Infection (BTI), BF.7 BTI, BA.1 BTI with reinfection, and BA.2 BTI with reinfection. Epitope groups predominantly comprising non-neutralizing or weakly neutralizing mAbs (E2.2, E3, and F1) are highlighted with dashed boxes. The percentage of antibodies in these three groups is labeled on each bar. **d**, Average DMS escape scores of the crucial epitope groups contributing to neutralization against XBB.1.5 are illustrated on the structure model of the SARS-CoV-2 BA.5 RBD (PDB: 7XNS). Key residues with high escape scores for each group are labeled. **e**, The average DMS escape scores for the key epitope groups are represented as sequence logos; residues are depicted using the standard one-letter code and colored based on their chemical properties. The height of each letter corresponds to the escape score of the respective mutation. **f**, Pseudovirus-neutralization activities of mAbs within the six crucial epitope groups (A1 [n = 170], A2 [n = 60], B [n = 33], F3 [n = 129], D3 [n = 155], and D4 [n = 80]) are shown against SARS-CoV-2 D614G, BA.5, BQ.1.1, and XBB.1.5. Geometric mean IC50 values are displayed as bars and labeled above each group of data points.

Neutralization data from both mice and human studies underscore the crucial role of secondary Omicron exposure in mitigating immune imprinting and generating potent antibody responses to immune-evasive variants such as XBB and its sublineages. We propose that this is largely attributable to the further expansion of Omicron-specific memory B cells *de novo* generated by the first Omicron exposure. To assess this hypothesis, we first analyzed the Omicron specificity of RBD-specific memory B cells from BTIs, BTIs+reinfection, and vaccine-naive reinfection cohorts through fluorescence-activated cell sorting (FACS). As we previously reported, in one-time Omicron BTI cohorts, more than 70% of the RBD-binding memory B cells also bound to WT, indicating that post-vaccination Omicron infection mainly recalls cross-reactive memory B cells elicited by WT-based vaccination, but rarely contains BA.1/BA.2-specific B cells (Fig. 3a). Subsequently, following an extended duration of time (8 months) after the first Omicron BTI, the proportion of cross-reactive cells declined while that of Omicron-specific cells increased, suggesting that longer B cell maturation periods elevated the proportion of Omicron-specific memory B cells (Fig. 3b). Nevertheless, at 8 months post-BA.1 BTI, the plasma neutralizing titers were very low due to antibody waning, and thus required a secondary Omicron boosting via either vaccination or infection to increase the antibody levels (Extended Data Fig. 3d). Importantly, for Omicron BTI+reinfection cohorts, the proportion of cross-reactive cells declined further but still remained higher than that observed in the vaccination-naïve reinfection cohort (Fig. 3c-d). These results are highly correlated with the plasma NT50s of the cohorts, which suggests that Omicron-specific antibodies are a major contributor for the increased antibody breadth and neutralization capability after repeated Omicron infection.

To further investigate the potency, breadth, and epitopes of these antibodies, the BA.1 RBD-binding cells and BA.2 RBD-binding cells from above various BA.1/BA.2 infection cohorts were sorted and sequenced by high throughput single-cell V(D)J sequencing. Antibodies were then expressed *in vitro* as human IgG1 monoclonal antibodies (mAbs) (Supplementary Table 2). For one-time Omicron BTI cohorts, enzyme-linked immunosorbent assay (ELISA) confirmed that only approximately 20% of the isolated mAbs specifically bind to the BA.1/BA.2 RBD and were not cross-reactive to the WT RBD, which was consistent with FACS results (Fig. 3e). Furthermore, long-term sampling (8 months) after BA.1 BTI yielded an increased proportion of BA.1 RBD-specific mAbs compared to short-term (2 months) sampling. Moreover, reinfection with BA.5/BF.7 further increased the proportion of BA.1/BA.2 RBD-specific mAbs to around 50%, but this was still lower than that in vaccination-naïve reinfection groups (Fig. 3e). Notably, the somatic hypermutation (SHM) rates of BA.1/BA.2 specific antibodies in BTI+reinfection cohorts were higher than that in one-time BTI cohorts (Fig. 3f), and the increased affinity maturation of BA.1/BA.2-specific antibodies contributes to their increased potency against Omicron variants (Fig. 3g-h). Together, these data indicate that long-term maturation after one-time Omicron BTI and repeated Omicron infections could significantly raise the proportion and maturation of Omicron-specific antibodies, greatly contributing to the increased plasma neutralization potency against Omicron variants.

### Epitope analyses of Omicron-specific mAbs

To further interrogate the composition of antibodies elicited by Omicron BA.5/BF.7 BTI and reinfection, and deciphering the molecular mechanism behind the broadly neutralizing capability of convalescent plasma from reinfection, we determined the binding sites and escaping mutations on RBD of these mAbs using deep mutational scanning (DMS)^39,40^. As the proportion of Omicron-specific antibodies is indispensable in reinfection cohorts, and the last exposure of all cohorts involved in this study is BA.5/BF.7, we built a yeast display mutant library based on BA.5 RBD and performed DMS for these mAbs in a high-throughput manner, akin to our previously described WT-based methods^40^. To enhance the sampling of Omicron-specific NAbs to facilitate the epitope characterization of these unprecedented antibodies, we specifically isolated an additional panel of RBD-targeting mAbs that do not cross-bind to WT according to the feature barcode counting during the 10x VDJ sequencing and determined their BA.5-based DMS data. We also determined the BA.5-based DMS data for all BA.5-RBD binding mAbs from previous collections isolated from various immune backgrounds (Supplementary Table 2). In total, a comprehensive panel consisting of 1350 mAb BA.5-based DMS is collected.

By graph-based unsupervised clustering on the determined escape scores over sites on RBD, we identified 12 major epitope groups on BA.5 RBD and embedded the mAbs using UMAP for visualization (Fig. 4a). Names of the epitope groups are generally assigned in line with the epitope groups on WT RBD defined previously^7,32^. Neutralizing activities against SARS-CoV-2 D614G, BA.1, BA.2, BA.5, BA.2.75, BQ.1.1, and XBB.1.5 are determined using VSV-based pseudovirus neutralization assays. In general, neutralization is highly correlated with targeting epitopes of mAbs. Antibodies in epitope groups F3, A1, A2, B, C/D1, D2, D3, D4, and E1/E2.1 target neutralizing epitopes, while antibodies in the other three groups, E2.2, E3, F1, exhibit weak or no neutralization activity (Fig. 4b and Extended Data Fig. 5b). Consistent with the plasma neutralization results, BA.5 or BF.7 BTI exhibited substantially imprinted antibody response, leading to over 50% antibodies that target conserved weakly neutralizing epitopes. In contrast, convalescent individuals who experienced BA.5 or BF.7 reinfection after prior BA.1 or BA.2 BTI induce only ∼20% antibodies targeting such epitopes, indicating striking alleviation of immune imprinting (Fig. 4c and Extended Data Fig. 5a). Interestingly, prior BA.1 or BA.2 BTI leads to Omicron-specific antibodies targeting distinct epitopes after reinfection. Prior BA.1 BTI induces a higher level of Group D3, while BA.2 BTI cohorts consist of more antibodies in Group F3, indicating that the Omicron infection history during repeated Omicron infections would also introduce new Omicron-based immune imprinting.

**Fig. 5.**
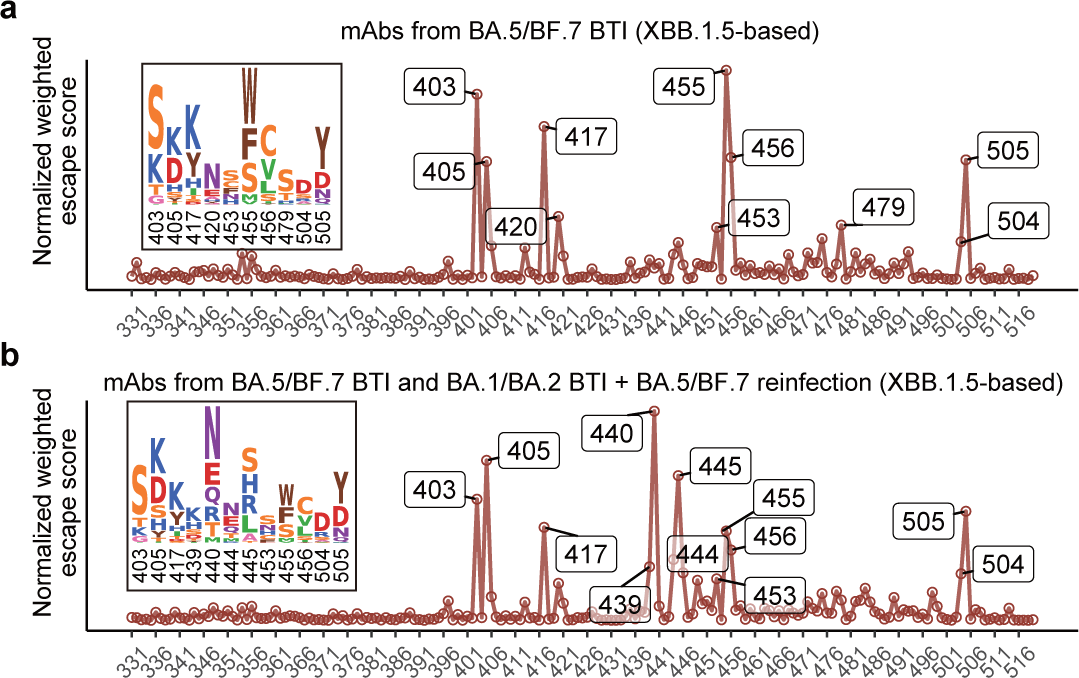
Estimate the evolutionary trends of XBB.1.5 RBD from DMS profiles. Normalized average DMS escape scores weighted by IC50 against XBB.1.5 using DMS profiles of mAbs from BA.5/BF.7 BTI (**a**), and mAbs from BA.5/BF.7 BTI and BA.1/BA.2 BTI with BA.5/BF.7 reinfection (**b**). The impacts of each mutation on ACE2 binding and RBD expression, and the codon constraints on each residue, are also considered (see Methods). Residues with high estimated preferences are labeled, and their corresponding mutation scores are shown as logos.

Among the 12 identified epitope groups, A1, D2, E1/E2.1, E2.2, E3, and F1 are similar to their corresponding WT-based groups and mainly consist of WT-reactive antibodies (Fig. 4d-e and Extended Data Fig. 5c-d)^32,41^. As expected, BA.5-based epitope landscape also defines novel groups that mainly comprise Omicron-specific mAbs, including Group A2, D3, D4, and F3. Notably, most antibodies in Group F3 here are not cross-reactive to WT RBD as well, which is different from the rare sarbecovirus-neutralizing broad NAbs in Group F3 from SARS convalescents described previously, such as SA55 and BD55-3372^42^. Compared to A1, which mainly contains IGHV3-53/3-66 public antibodies (also known as class 1 or site Ia)^43,44^, mAbs in Group A2 are susceptible to mutations on 417 and 505, including the reversions. Group D3 and D4 target an epitope near Group D2 (targeted by LY-CoV1404), but exhibited distinct escape profiles or interacting residues^45^. D3 is susceptible to N439 and K440 mutations, and thus escaped by WT due to N440, while the footprint of D4 is closer to the receptor-binding motif (RBM), interacting with G447, Y449, and R498 (Fig. 4d-e). Antibodies in WT-based Group B, C, and D1 have been mostly escaped by L452R, E484A, and F486V in BA.5. B and C/D1 here comprise both WT-reactive and Omicron-specific antibodies, where Group B is more focused on N487 and Y489, and C/D1 mainly focus on F490, which is largely escaped by F490S in XBB variants (Fig. 4d-e and Extended Data Fig. 5c-d). Among the 12 groups, A1, A2, B, D3, especially D4 and F3 consist of a substantial proportion of NAbs exhibiting broad neutralization against BQ.1.1 and XBB.1.5 (Fig. 4f). C/D1, D2, and E1/E2.1 also consist of a small proportion of XBB.1.5-neutralizing mAbs (Extended Data Fig. 5f). Considering the recent emergence and prevalence of XBB subvariants harboring F456L (XBB.1.5.10) or K478R (XBB.1.16), which are crucial sites for NAbs in Group A1 and A2, or B and C/D1, respectively, we tested the neutralization of XBB.1.5-neutralizing antibodies from these groups against these two mutants. As expected, F456L escapes or dampens the neutralization of most XBB.1.5-neutralizing antibodies in Group A1 or A2, and XBB.1.16 (E180V+K478R) also escapes a large proportion of NAbs in B and C/D1 (Extended Data Fig. 5e). Overall, these results demonstrate that Omicron repeated infection stimulates a higher level of Omicron-specific neutralizing antibodies targeting neutralizing epitopes compared to one-time Omicron BTI, indicating substantial alleviation of immune imprinted on antibody epitope level. And that these Omicron-specific mAbs have distinct RBD epitopes and escaping mutations compared to WT-induced mAbs would introduce a large neutralizing epitope shift, contributing majorly to the broadly neutralizing capability against XBB.1.5.

### Evolutionary hotspots on XBB.1.5 RBD

Encouraged by the successful rationalization of the prevalence of F456L and K478R based on DMS, we desire to systematically investigate the evolutionary preference of other RBD mutations. To integratively evaluate the preference of each mutation considering their impacts on neutralizing antibody escape, hACE2 binding, RBD stability, and codon constraints, we previously calculated a weighted preference score for RBD mutations using WT-based DMS profiles and neutralizing activities against BA.5 to predict the convergent evolution of BA.5 RBD^7^ (Extended Data Fig. 6). We desire to utilize similar approach with BA.5-based profiles and neutralization against XBB.1.5 to identify the evolutionary trends of XBB.1.5 RBD. When considering antibodies from BA.5/BF.7 BTI only, the most significant sites include R403S/K, N405K, N417Y, Y453S/C/F, L455W/F/S, F456C/V/L, and H505Y/D, corresponding to escape hotspots of Group A1, A2, and F3 (Fig. 5a). With antibodies from repeated Omicron infection included in the analysis, scores of N439K, K440N/E, K444N/E, and P445S/H/R/L become higher, corresponding to Group D3 and D4, which are consistent with the epitope distributions of mAbs from each cohort (Fig. 5b). Notably, N405D and N417K reversions should hardly appear in the real world due to the potential recovery of previously escaped NAbs in Group F2 and A, respectively. K478 mutations are not identified in the calculation, which is also a limitation of our model due to the low proportion of XBB-neutralizing antibodies in Group B or C/D1 in our cohorts.

**Fig. 6.**
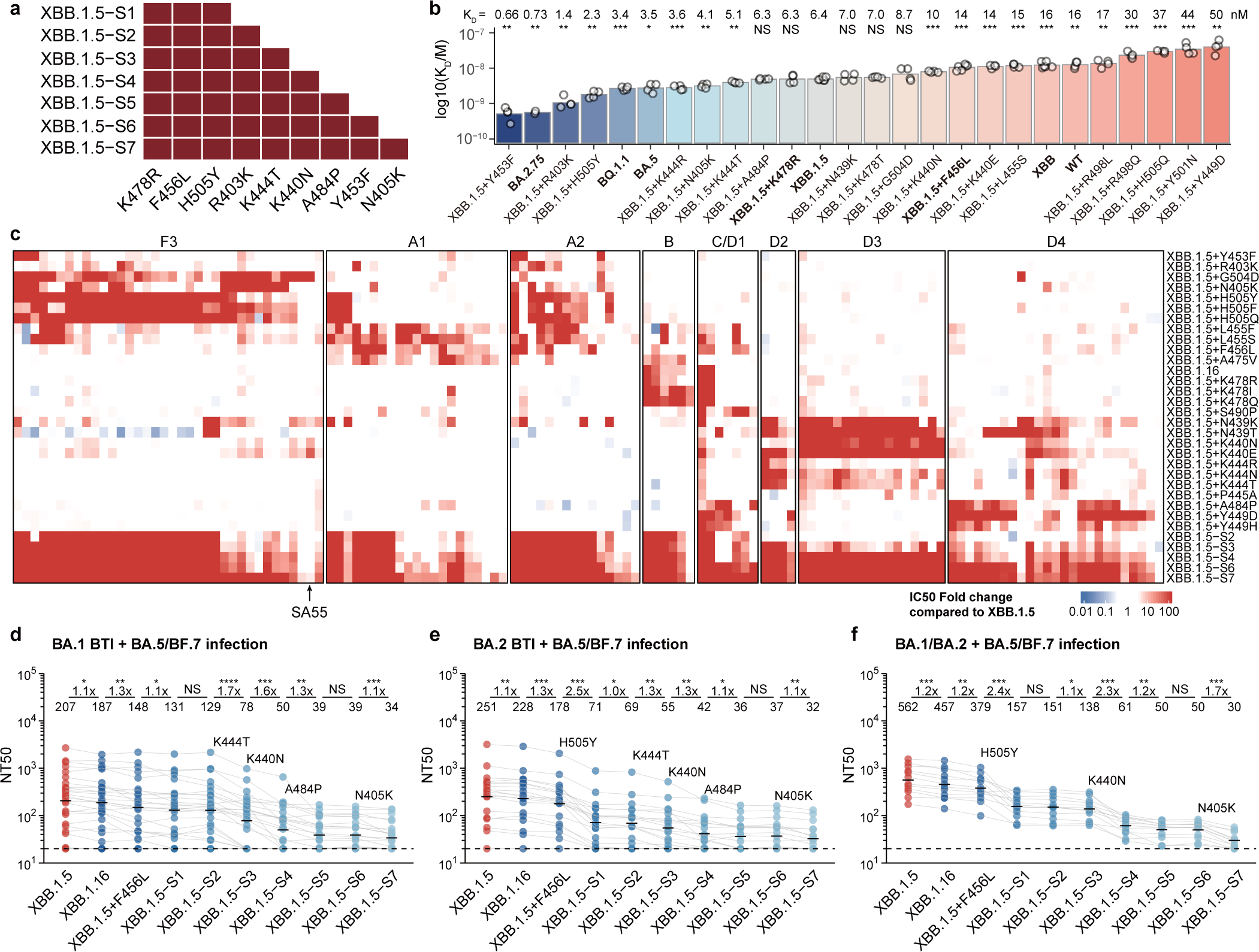
Combination of escape mutations evades XBB.1.5-neutralizing antibodies from reinfection. **a**, SARS-CoV-2 XBB.1.5-based pseudoviruses harboring combinations of critical mutations identified through analysis of DMS profiles are generated. **b**, hACE2-binding affinity for various RBD mutants of SARS-CoV-2 is assessed using SPR. Geometric mean dissociation constants (KD) from at least four independent replicates are shown, with statistical significance in comparison to XBB.1.5 RBD’s KD labeled above the bars. P-values are determined using a two-tailed t-test on log-transformed KD values. **c**, IC50 values for representative potent XBB.1.5-neutralizing antibodies from different epitope groups against XBB.1.5 variants carrying individual or multiple escape mutations are displayed. Fold changes in IC50 against the mutants relative to XBB.1.5 are presented as a heatmap. **d-f**, Pseudovirus 50% neutralization titers (NT50) for SARS-CoV-2 XBB.1.5-based mutants are shown using plasma from convalescent individuals who experienced BA.5 or BF.7 reinfection: BA.1 BTI prior to BA.5/BF.7 reinfection (n = 26) (**d**); BA.2 BTI prior to BA.5/BF.7 reinfection (n = 19) (**e**); and reinfection with BA.5 or BF.7 after BA.1 or BA.2 infection without vaccination (n = 12) (**f**). Key mutations diminishing neutralization are labeled above their corresponding lines. Dashed lines indicate the limit of detection (LOD, NT50 = 20). Geometric mean titers are labeled above data points. Statistical tests are performed between neighboring mutants. P-values are calculated using two-tailed Wilcoxon signed-rank tests on paired samples. *p < 0.05, **p < 0.01, ***p<0.001, ****p<0.0001, and not significant (NS) p > 0.05.

Based on the analysis above, we wonder if the combination of multiple escape mutations against major XBB.1.5-effective epitope groups could essentially evade the broadly neutralizing capability of plasma from repeated Omicron infection while retaining high ACE2 binding affinity. Besides the two emerging mutations K478R and F456L, we selected seven additional substitutions, including H505Y, R403K, K444T, K440N, A484P, Y453F, and N405K, which are sequentially added to XBB.1.5, and constructed seven pseudoviruses named XBB.1.5-S1 to XBB.1.5-S7 (Fig. 6a). The mutations are selected from a larger set of mutation candidates considering their impacts on hACE2-binding affinity as determined by surface plasmon resonance (SPR) and the capability of escaping the neutralization of a panel of 131 potent XBB.1.5-neutralizing antibodies from 8 epitope groups (Fig. 6b-c and Extended Data Fig. 7a). XBB.1.5-S7 successfully escapes the most of NAbs in the panel, except for a small group of broad NAbs from Group F3, A1, and D4, including SA55, a therapeutic antibody under clinical development^42^. Then, we evaluated the neutralization titers of convalescent plasma from individuals who experienced Omicron BTI or repeated Omicron infection against the designed escape mutants. As expected, XBB.1.5-S7 could significantly escape plasma samples from all tested cohorts. Plasma from BA.5 or BF.7 BTI are significantly escaped upon the inclusion of F456L, and nearly negative against XBB.1.5-S7 (Extended Data Fig. 7b). Plasma from repeated Omicron infections is much more resistant to escape mutations. Interestingly, plasma from BA.5/BF.7 reinfection with prior BA.1 BTI or BA.2 BTI exhibited distinct neutralization to different escape mutants. The former samples are largely evaded by K444T and K440N, but not strongly affected by H505Y, while the latter samples are significantly evaded by H505Y (Fig. 6d-e). This is consistent with the observation that reinfection with prior BA.1 BTI elicits more Group D3 antibodies, while reinfection with prior BA.2 BTI elicits more Group F3 antibodies (Fig. 4c). Unvaccinated reinfection cohorts exhibited higher neutralization against XBB.1.5 compared to vaccinated cohorts, but equivalently escaped by XBB.1.5-S7. The most significant reduction occurs upon the inclusion of H505Y, K440N, and N405K, indicating a high proportion of Omicron-specific antibodies in Group D3 and F3 (Fig. 6f).

In summary, our findings suggest that secondary Omicron exposure is necessary to mitigate the immune imprinting conferred by previous ancestral virus exposure and to elicit higher levels of Omicron-specific antibodies. Accordingly, our recommendation is to administer two booster doses of Omicron-based vaccines to individuals who have not received prior Omicron-based vaccinations or who have not been previously infected with the Omicron variant. Moreover, administering the second booster shot after a prolonged interval can provoke a wider and more efficient immune response, while incorporating the wildtype virus into subsequent vaccine designs may worsen immune imprinting^26^. Furthermore, it is imperative to incorporate the XBB variants into vaccine design to achieve broad-spectrum protection, given its potential to mutate and evade vaccines based on previous Omicron variants.

Recently, several fast-growing XBB lineages, such as the variant of interest (VOI) XBB.1.16 (K478R), XBB.2.3.5 (K478N), and XBB.2.3.4 (K478Q), have acquired RBD mutations on K478. However, the K478 mutation did not emerge in our prediction of evolutionary trends for XBB.1.5 RBD. This contradiction may be attributed to the fact that our mutational prediction model primarily relies on the cohorts we recruited, and we haven’t captured the immune background that introduced K478 mutation. One possible background that may give rise to K478 is repeated BA.5/BQ.1.1/XBB exposure, as F486 could mask the immunogenicity of K478. Another potential source of K478 is Delta-imprinted convalescents who experienced BA.5/BQ.1.1/XBB infections, which could result in the generation of abundant K478X-sensitive mAbs, given that Delta carries T478K. This may explain why K478X is mostly observed in India^4,46^.

The degree of immune imprinting might be different between mRNA and inactivated vaccination. Recent studies have shown that subsequently exposed to Omicron twice after two doses of WT-based mRNA vaccines still produce significantly low levels of Omicron-specific antibodies, despite the enhanced neutralization breadth against BQ.1.1 and XBB variants^47,48^. Additionally, individuals who have received two doses of mRNA vaccines and experienced two rounds of Omicron infection also have low levels of Omicron-specific antibodies^47^. This indicates that mRNA vaccines may generate a stronger immune imprinting effect compared to inactivated vaccines, potentially due to its stronger primary humoral immune response^8,49^. However, a head-to-head comparison is needed for validation.

## Methods

### Isolation of PBMCs and plasma

Blood samples from vaccinated or unvaccinated individuals who had recovered from Omicron breakthrough infection or reinfection were obtained under study protocols approved by Beijing Ditan Hospital, Capital Medical University (Ethics committee archiving No. LL-2021-024-02) and the Tianjin Municipal Health Commission, and the Ethics Committee of Tianjin First Central Hospital (Ethics committee archiving No. 2022N045KY). All participants have provided written informed consent for the collection of information, storage and use of their clinical samples for research purposes, and publication of data generated from this study.

Samples from one-time breakthrough infection and the first infections in repeat-infection cohorts were collected during the “zero COVID” period in China. During that period, the total number of infected individuals was small and there were clear epidemiological correlations between confirmed cases. BA.1 breakthrough infections occurred in Tianjin in January and a cumulative count of 430 individuals tested positive for Omicron BA.1 by February 7, 2022, with no additional infections identified in the subsequent 16 days ^38^. BA.2 breakthrough infections occurred in Beijing between April and July 2022. From April 22 to Nov 14, a total of 2,230 cases of local infections were reported in Beijing, and BA.2.2.1 (BA.2+I1221T in spike) was the most prevalent subvariant in Beijing between April and July^50^. BA.5 breakthrough infections occurred in Beijing and Tianjin between September and October 2022^50^. BF.7 breathrough infections occurred in Inner Mongolia in November 2022, and BF.7 accounted for 100% of the sequences^51^. These samples of infection were confirmed by PCR, and the majority of them also underwent sequencing to determine the viral strains. The unsequenced samples, which make up only a small proportion of the total samples, showed strong epidemiological correlations with the sequenced samples.

Reinfections were confirmed by PCR or antigen testing. While the viral strain types for these infections were not confirmed through sequencing, it is important to note that these samples were confirmed in December 2022 in Beijing and Tianjin. At that time, these regions were predominantly undergoing the BA.5/BF.7 wave^50^. Among the sequences from samples collected between 12/01/2022-02/01/2023, >98% of them were designated as BA.5* (excluding BQ*). Specifically, the major subtypes circulating in China at that time were BA.5.2.48* (DY*) and BF.7.14*, which do not harbor additional mutations on RBD, and thus can be generally considered as BA.5/BF.7 in this study (https://cov-spectrum.org/explore/China/AllSamples/from%3D2022-12-01%26to%3D2023-02-01/variants?&).

The whole blood samples were 1:1 diluted with 2% fetal bovine serum (FBS) (Hyclone, SH30406.05) in phosphate buffered saline (PBS) (Invitrogen, C10010500BT) and subjected to Ficoll (Cytiva, 17-1440-03) gradient centrifugation to isolate plasma and PBMCs. Plasma was collected from upper layer after centrifugation. PBMCs were collected at the interface and further prepared through centrifugation, red blood cell lysis (Invitrogen™ eBioscience™ 1X RBC Lysis Buffer, 00-4333-57) and washing steps. If not used for downstream process immediately, samples were stored in FBS with 10% DMSO (Sigma-Aldrich, D4540) in liquid nitrogen. All PBMC samples were shipped on dry ice and cryopreserved PBMCs were thawed in PBS + 1mM EDTA (Invitrogen, AM9260G) + 2% FBS before use.

### mRNA and protein vaccine preparation and mouse immunization

For mRNA vaccine preparation, 5′ untranslated region (UTR), target sequence, and 3’UTR were sequentially inserted after T7 promoter in an empty PSP73 plasmid firstly. The plasmid was then subjected to double digestion to obtain linearized DNA. This DNA served as a template for an in vitro transcription reaction mediated by T7 RNA polymerase to synthesize RNA encoding the SARS-CoV-2 S6P (F817P, A892P, A899P, A942P, K986P, V987P, R683A and R685A) protein according to the manufacturer’s instructions (Vazyme, DD4201). Uridine was fully replaced by N1-methyl-pseudouridine in this process. Transcription products were treated with DNase I to remove DNA templates, and purified using VAHTS RNA Clean Beads (Vazyme, N412-02). Cap 1 structure was added using Vaccinia Capping Enzyme (Vazyme, DD4109) and mRNA Cap 2’-O-Methyltransferase (Vazyme, DD4110), followed by magnetic bead purification. Poly(A) tails were added using E.coli Poly(A) Polymerase (Vazyme, N412-02) and the product was purified again.

The mRNA was encapsulated in a functionalized lipid nanoparticle as described previously^52^. In brief, ionizable lipid, DSPC, cholesterol, and PEG2000-DMG were dissolved in ethanol at the mole ratio of 50:10:38.5:1.5, respectively. mRNA was diluted in RNase free 50 mM citrate buffer (pH 4.0) to obtain a final lipid:mRNA weight ratio of 6:1. The aqueous and ethanol solutions were mixed in a 3:1 volume ratio using a microfluidic apparatus and the obtained LNPs were dialyzed overnight. All of the samples were stored within a week at 2∼8 ℃ of use to ensure the chemical stability of the components. The size of LNPs, the particle size distributions, and the encapsulation and concentration of mRNA were determined. The encapsulation in all of the samples was typically 90– 99%.

The spike proteins, including D614G (ACROBiosystems, SPN-C52H9), XBB (ACROBiosystems, SPN-C5248), BQ.1.1 (ACROBiosystems, SPN-C522s), BA.1 (ACROBiosystems, SPN-C522a), BA.5 (ACROBiosystems, SPN-C522e) were used for mouse immunization. All of these proteins were modified to incorporate 6P2A mutations (F817P, A892P, A899P, A942P, K986P, V987P, R683A, R685A) and a T4 fibritin foldon domain at the C-terminus to improve the stability of the trimeric structure.

Animal experiments were carried out under study protocols approved by Institute of Biophysics, Chinese Academy of Sciences (SYXK2023300) and HFK Biologics (HFK-AP-20210930). Mice were immunized according to schemes in figure 1. All inactivated vaccines were administered intraperitoneally at a dose of 3 μg per mouse, while mRNA vaccines were administered intramuscularly at a dose of 10 μg per mouse. Protein subunit vaccines were administered subcutaneously at six sites on the back at a dose of 10 μg per mouse, where complete Freund’s adjuvant was used for the prime immunization, and incomplete Freund’s adjuvant was used for booster immunizations, with a 1:1 volume ratio of protein subunit and adjuvant. The second immunizations were given 2 weeks after the first dose, with subsequent doses administered at 1-month intervals, unless stated otherwise. Blood samples were collected 1 week after the final immunization.

### BCR sequencing, analysis and recombinant antibody expression

CD19^+^ B cells were enriched from PBMCs using EasySep Human CD19 Positive Selection Kit II (STEMCELL, 17854). Following enrichment, 1x10^6^ B cells in 100 μl buffer were incubated with a panel of antibodies including 3 μl FITC anti-human CD20 antibody (BioLegend, 302304), 3.5 μl Brilliant Violet 421 anti-human CD27 antibody (BioLegend, 302824), 2 μl PE/Cyanine7 anti-human IgD antibody (BioLegend, 348210) and 2 μl PE/Cyanine7 anti-human IgM antibody (BioLegend, 314532). Additionally, fluorophore or oligonucleotide conjugated RBD were added. For FACS, 0.013 μg of biotinylated BA.1 (Sino Biological, 40592-V49H7-B) or BA.2 (customized from Sino Biological) RBD protein conjugated with PE-streptavidin (BioLegend, 405204) and APC-streptavidin (BioLegend, 405207), and 0.013 μg of WT biotinylated RBD protein (Sino Biological, 40592-V27H-B) conjugated with BV605-streptavidin (BioLegend, 405229) were added. For sequencing, BA.1 or BA.2 biotinylated RBD protein conjugated with TotalSeq™-C0971 Streptavidin (BioLegend, 405271) and TotalSeq™-C0972 Streptavidin (BioLegend, 405273), WT biotinylated RBD protein conjugated with TotalSeq™-C0973 Streptavidin (BioLegend, 405275) and TotalSeq™-C0974 Streptavidin (BioLegend, 405277) and biotinylated Ovalbumin (Sino Biological) conjugated with TotalSeq™-C0975 Streptavidin (BioLegend, 405279) were added. After incubation and washing steps, 5 μl of 7-AAD (Invitrogen, 00-6993-50) was included for dead cell exclusion.

Cells negative for 7-AAD, IgM and IgD, but positive for CD20, CD27 and BA.1 or BA.2 were sorted using a MoFlo Astrios EQ Cell Sorter (Beckman Coulter). FACS data were collected by Summit 6.0 (Beckman Coulter) and analyzed using FlowJo v10.8 (BD Biosciences).

The sorted B cells were processed using the Chromium Next GEM Single Cell V(D)J Reagent Kits v1.1 according to the manufacturer’s user guide (10x Genomics, CG000208). Briefly, the cells were resuspended in PBS after centrifugation and then processed to obtain gel beads-in-emulsion (GEMs) using the 10X Chromium controller. The GEMs were subjected to reverse transcription and the products were further purified with a GEM-RT clean up procedure. Preamplification was then performed on the products which were subsequently purified using the SPRIselect Reagent Kit (Beckman Coulter, B23318). The paired V(D)J BCR sequences were enriched with 10X BCR primers, followed by library preparation. Finally, the libraries were sequenced using the Novaseq 6000 platform, running either the Novaseq 6000 S4 Reagent Kit v1.5300 cycles (Illumina, 20028312) or the NovaSeq XP 4-Lane Kit v1.5 (Illumina, 20043131).

10X Genomics V(D)J sequencing data were assembled as BCR contigs and aligned using the Cell Ranger (v6.1.1) pipeline according to the GRCh38 BCR reference. To ensure high quality, only the productive BCR contigs and cells with one heavy chain and one light chain were retained. The IgBlast program (v1.17.1) was utilized to identify and annotate the germline V(D)J genes. The Change-O toolkit (v1.2.0) was employed to detect somatic hypermutation sites in the variable domain of the antibodies.

For expression optimization in human cells, heavy and light chain genes were synthesized by GenScript, inserted separately into plasmids (pCMV3-CH, pCMV3-CL or pCMV3-CK) via infusion (Vazyme, C112), and co-transfected into Expi293F cells (Thermo Fisher, A14527) using polyethylenimine transfection. The cells were cultured at 36.5°C in 5% CO_2_ and 175 r.p.m. for 6-10 days. The cell expression fluid was collected and centrifuged. After centrifugation, supernatants containing the monoclonal antibodies were purified using protein A magnetic beads (Genscript, L00695). The purified samples were determined by SDS-PAGE.

### Pseudovirus-neutralization assay

Codon-optimized SARS-CoV-2 S gene was inserted into the pcDNA3.1 vector to construct plasmids encoding the spike proteins of SARS-CoV-2. The 293T cell line (ATCC, CRL-3216) was transfected with the spike protein-expressing plasmids and then infected with G*ΔG-VSV virus (Kerafast, EH1020-PM). After culturing, the pseudovirus-containing supernatant was collected, filtered, aliquoted, and frozen at −80 °C for future use. Pseudovirus-neutralization assays were conducted on the Huh-7 cell line (Japanese Collection of Research Bioresources (JCRB), 0403).

Monoclonal antibodies or plasma were serially diluted in DMEM (Hyclone, SH30243.01) and incubated with pseudovirus in 96-well plates at 5% CO_2_ and 37°C for 1 h. Digested Huh-7 cell (JCRB, 0403) or 293T-hACE2 cells (ATCC, CRL-3216) were seeded and cultured for 24h. Half of the supernatant was then discarded and D-luciferin reagent (PerkinElmer, 6066769) was added to react in the dark. The luminescence value was detected using a microplate spectrophotometer (PerkinElmer, HH3400). IC50 was determined by a four-parameter logistic regression model using PRISM (version 9.0.1).

### Authentic virus neutralizing assay

The serum samples obtained from Convalescent individuals were heat-inactivated at 56°C for 0.5 hours and subsequently diluted in two-fold steps with cell culture medium. These diluted sera were mixed with a virus suspension (SARS-CoV-2 Wuhan, BA.1, BA.5.2.1, BF.7.14, FL.8 (XBB.1.9.1.8) containing 100 CCID_50_ and added to 96-well plates at a 1:1 ratio. The plates were then incubated at 36.5°C in a 5% CO_2_ incubator for 2 hours. Following the incubation period, Vero cells (Gifted from WHO, (ATCC, CCL-81))were added to each well containing the serum-virus mixture. The plates were further incubated for 5 days at 36.5°C in a 5% CO_2_ incubator. Microscopic observation of cytopathic effects (CPE) was performed, and the neutralizing titer was determined based on the highest dilution that showed 50% protection against the virus-induced CPE.

### ELISA

ELISA assays were conducted by pre-coating ELISA plates with RBD (SARS-CoV-2 wild type, SARS-CoV-2 BA.1, SARS-CoV-2 BA.2 RBD, Sino Biological) at concentrations of 0.03 μg ml^−1^ and 1 μg ml^−1^ in PBS overnight at 4 °C. The plates were then washed and blocked, after which 100 μl of 1 μg ml^−1^ antibodies were added to each well and incubated at room temperature for 2 hours. Following incubation, the plates were washed and incubated with 0.25 μg ml^−1^ Peroxidase-conjugated AffiniPure goat anti-human IgG (H+L) (JACKSON, 109-035-003) for 1 hour at room temperature. The reaction was developed using tetramethylbenzidine (TMB) (Solarbio, 54827-17-7), and stopped by adding H_2_SO_4_. The absorbance was measured at 450 nm using a microplate reader (PerkinElmer, HH3400) and the negative control used was the H7N9 human IgG1 antibody HG1K. (Sino Biological, HG1K).

### Surface plasmon resonance

Human ACE2 with Fc tag was immobilized onto Protein A sensor chips using a Biacore 8K (GE Healthcare). Purified SARS-CoV-2 mutants RBD were prepared in serial dilutions, ranging from 100 to 6.25 nM, and injected over the sensor chips. The response units were recorded at room temperature using BIAcore 8K Evaluation Software (v3.0.12.15655; GE Healthcare). The obtained data were then analyzed using BIAcore 8K Evaluation Software (v3.0.12.15655; GE Healthcare) and fitted to a 1:1 binding model.

### DMS Library construction

Duplicate single site saturated mutant libraries spanning all 201 amino acids of BA.5 RBD (position N331-T531 by Wuhan-Hu-1 reference numbering) were constructed based on previously reported method^1^, in order to ensure the reproducibility and reliability of results. A unique N26 barcode was PCR appended to each RBD variant as an identifier, and the correspondence of variant and N26 barcode was obtained by PacBio sequencing on Sequel ll platform in Peking University throughput sequencing center (HTSC). The BA.5 RBD mutant libraries were assembled into pETcon 2649 vector and amplified in DH10B cells. Above plasmids products were then transformed into Saccharomyces cerevisiae EBY100. Yeasts were screened on SD-CAA plates and further enlarged in SD-CAA liquid media, the resulted libraries were preserved at -80°C after flash frozen in liquid nitrogen.

### MACS-based mutation escape profiling

The high-throughput mutation escape profiling for every single antibody was performed as previously described^7,32^. Briefly, unexpressed and non-functional RBD variants were first eliminated from BA.5 mutant libraries by magnetic-activated cell sorting (MACS). The selected yeasts were inoculated into SG-CAA media to induce RBD surface expression by overnight culture. To capture yeast cells that escape specific antibody binding, two rounds of sequential negative selection and one round of positive selection were carried out based on MACS. After overnight amplification, plasmids were extracted from the sorted yeasts using the 96 Well Plate Yeast Plasmid Preps Kit (Coolaber, PE053), then used as template for N26 barcode amplification by PCR. Final PCR products were purified, quantified, and sequenced on Nextseq 550 or MGISEQ-2000 platform.

### DMS data analysis and antibody clustering

DMS raw sequencing data were processed as described previously^7,32^. In brief, the detected barcode sequences of both the antibody-screened and reference library were aligned to the barcode-variant dictionary generated using dms_variants (v0.8.9) from PacBio sequencing data of the BA.5 DMS library. Only barcodes that are detected more than 5 times in the reference library are included in the calculation to avoid large sampling error. The escape scores of a variant X that are detected both in the screened and reference library were defined as F×(n_X,ab_ / N_ab_) / (n_X,ref_ / N_ref_), where F is a scale factor to normalize the scores to the 0-1 range, while n and N are the number of detected barcodes for variant X and total barcodes in antibody-screened (ab) or reference (ref) samples, respectively. To assign an escape score to each single substitution on RBD, an epistasis model is fitted using dms_variants (v0.8.9) as described previously^53,54^. For antibodies with multiple replicates of DMS, the final escape score of each mutation is the average over all replicates.

We used graph-based unsupervised clustering and embedding to assign an epitope group for each antibody and visualize them in a two-dimensional space. First, site escape scores (the sum of mutation escape scores on a residue) of each antibody are first normalized to a sum of one and considered as a distribution over RBD residues. The dissimilarity of two antibodies is defined by the Jessen-Shannon divergence of the normalized escape scores. Pair-wise dissimilarities of all antibodies in the dataset are calculated using the SciPy module (scipy.spatial.distance.jensenshannon, v1.7.0). Then, a 12-nearest-neighbor graph is built using python-igraph module (v0.9.6). Leiden clustering is performed to assign a cluster to each antibody ^55^. The name of each cluster is annotated manually based on the featured sites on the average escape profiles of a cluster to make it consistent with the definition of our previously published DMS dataset using WT-based library in general^7^. To project the dataset onto a 2D space for visualization, we performed UMAP based on the constructed k-nearest-neighbor graph using umap-learn module (v0.5.2). Figures were generated by R package ggplot2 (v3.3.3).

### Estimate the preference of RBD mutations

Similar to the approach in our previous study^7^, we incorporated four types of weights in our calculations to account for the impact of each mutation on hACE2-binding affinity, RBD expression, neutralizing activity, and the codon constraints on each residue. The weights for ACE2 binding and RBD expression are determined by tanh(*S*_bind_) + 1 and tanh(min(0, *S*_expr_)) + 1, respectively, where the S_bind_ and S_expr_ values are from the BA.2-based DMS on ACE2 binding and RBD expression^56^. The function tanh(*x*) is employed as a sigmoidal curve to constrain the weights between 0 and 2. For codon constraint weights, mutations that cannot be accessed through single nucleotide mutation are first assigned a weight of zero. To address the intrinsic disparities in the frequency of distinct nucleotide substitutions in SARS-CoV-2, we assign different weights for mutations corresponding to various nucleotide substitutions^57^. Specifically, the weight of the most frequent substitution (C>T) is assigned a value of 0.1, while weights for G>T and G>A are 0.041 and 0.035, respectively. To retain the potential of rare mutations, all other substitutions are assigned a weight of 0.03. We use BA.4/5 (EPI_ISL_11207535) and XBB.1.5 (EPI_ISL_17054053) to define weights for codon usage. Regarding the neutralizing activities, the weight is calculated as - log_10_(IC_50_). IC_50_ values (μg/mL) less than 0.0005 or greater than 1.0 are considered as 0.0005 or 1.0, respectively. As the dataset specifically enriches for Omicron-specific antibodies, potentially introducing bias when estimating mutation preferences. An additional weighting strategy is applied that assigns higher weights to cross-reactive mAbs, resulting in 89% cross-reactive mAbs for BA.5/BF.7 BTI cohorts and 51% for reinfection cohorts, as determined by unbiased characterization of mAbs using ELISA. The raw escape scores for each antibody are first normalized by the maximum score among all mutants. The weighted score for each antibody and each mutation is obtained by multiplying the normalized scores with the corresponding four weights, and the final mutation-specific weighted score is the sum of scores for all antibodies in the designated set, which is then normalized once more to produce a value between 0 and 1. To visualize the calculated escape maps, sequence logos were created using the Python module logomaker (v0.8).

## Acknowledgments

We would like to express our gratitude to Jesse D. Bloom and Tyler N. Starr for their invaluable insights and discussions during the construction of our deep mutation scanning libraries. We thank all volunteers for providing the blood samples. We gratefully acknowledge the assistance of the High-throughput Sequencing Center at Peking University and Annoroad Gene Technology Co., Ltd. in sample sequencing. This project is financially supported by the Ministry of Science and Technology of China and Changping Laboratory (2021A0201; 2021D0102), and National Natural Science Foundation of China (32222030).

## Author contributions

Y.C. designed and supervised the study. A.Y., F.J., W.S., Q.G., X.S.X, and Y.C. wrote the manuscript with inputs from all authors. A.Y., W.S., S.Y., R.A., Yao W., and X.N. performed B-cell sorting, single-cell VDJ sequencing, and antibody sequence analyses. J.W. (BIOPIC), F.J., H.S. and L.Z. performed and analyzed the DMS data. Y.Y. and Youchun W. constructed the pseudotyped virus. N.Z., P.W., L.Y., T.X. and F.S. performed the pseudotyped virus neutralization assays, ELISA, and SPR. Z.L. performed authentic virus neutralization assays. W.S. and A.Y. analyzed the neutralization data. Y.X., X.C., Z.S. and R.J. recruited the SARS-CoV-2 vaccinees and convalescents. J.W. (Changping Laboratory), L.Y. and F.S. performed the antibody expression.

## Data and code availability

Information of SARS-CoV-2 RBD-targeting mAbs is included in Supplementary Table 2. Processed mutation escape scores, and custom scripts for processing and analyzing DMS data can be downloaded at https://github.com/jianfcpku/SARS-CoV-2-reinfection-DMS. Raw sequencing data of DMS are available on China National GeneBank with Project accession CNP0004294. PDB 7XNS is used for the structural model of SARS-CoV-2 RBD.

## Extended Data Figure Legends

**Extended Data Fig. 1.**
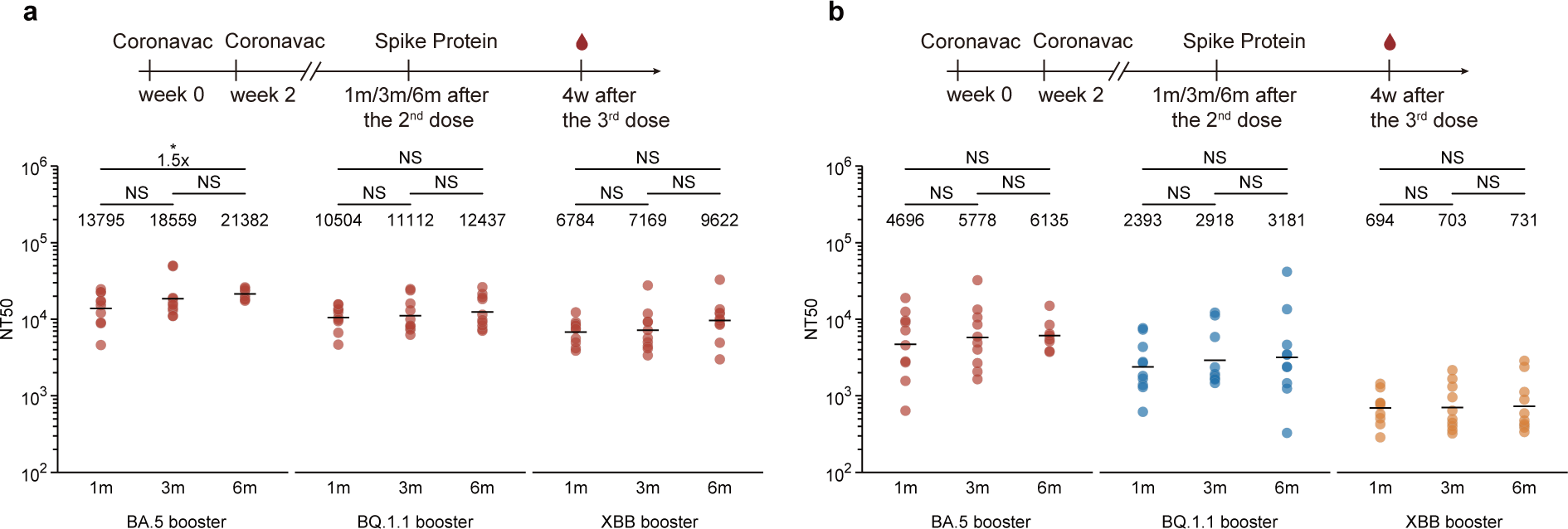
Neutralizing antibody response after CoronaVac priming and one-dose variant spike boosting. **a, b,** Comparison of neutralizing titers among different groups of mice immunized with 2 doses of CoronaVac followed by one-dose BA.5/BQ.1.1/XBB Spike protein boosters administered with one-month, three-month, or six-month intervals between the second and third dose. **a**) Neutralizing titers against D614G; **b**) Neutralizing titers against variants that the mice boosted with. Statistical significance was determined using the Wilcoxon rank sum test. *p < 0.05, **p < 0.01, ***p<0.001, ****p<0.0001, and not significant (NS) p > 0.05.

**Extended Data Fig. 2.**
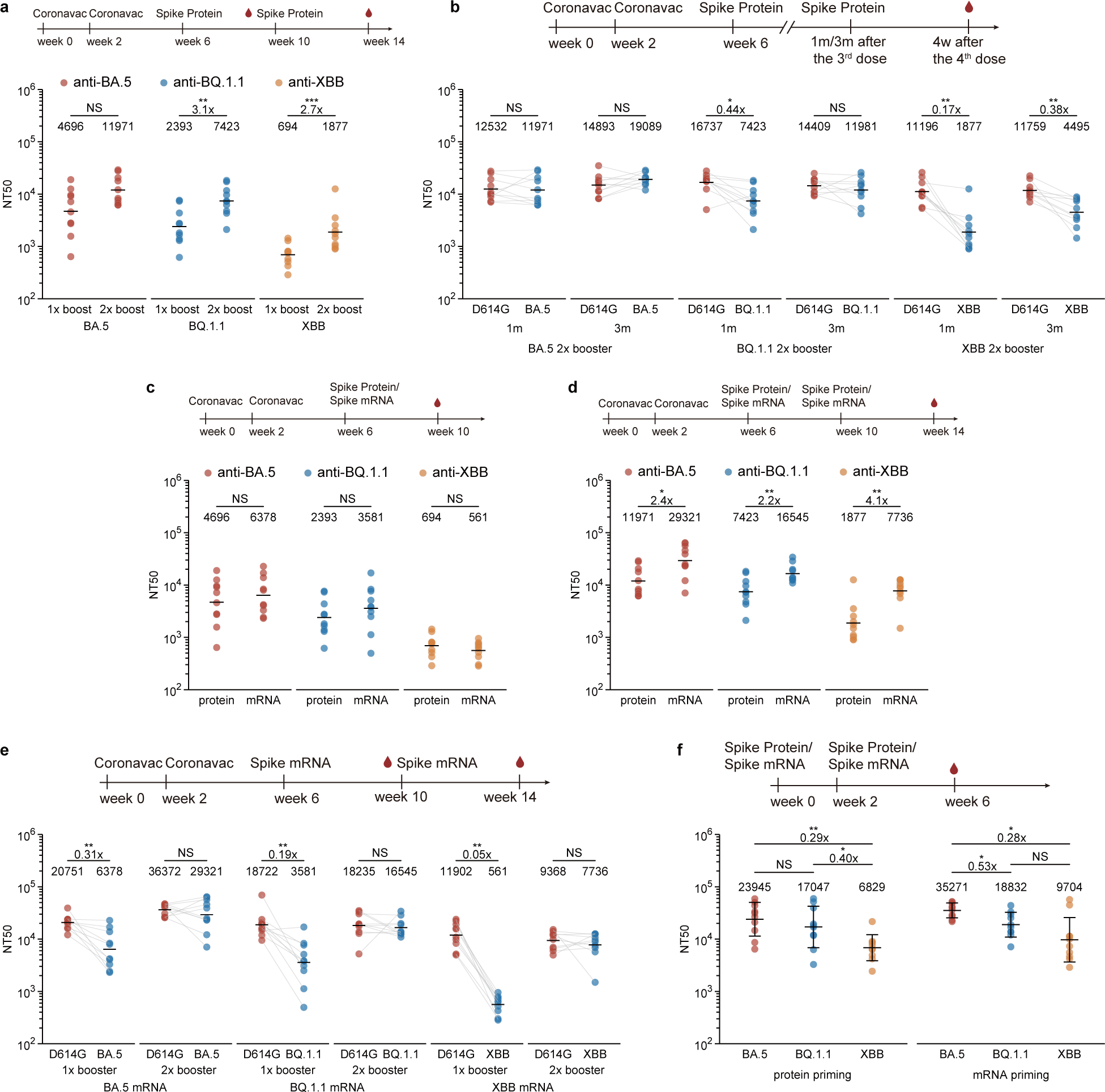
Neutralizing antibody response after CoronaVac priming and two-dose variant spike booster or two-dose variant spike priming. **a**, Comparison of neutralizing titers after CoronaVac priming and one-dose or two-dose variant spike boosting. **b**, D614G and boosting variant neutralizing titers after CoronaVac priming and two-dose variant spike boosting. **c-d**, Comparison of neutralizing titers after CoronaVac priming and variant spike protein or mRNA boosting. one-dose boosting in **c** and two-dose boosting in **d**. **e**, Neutralizing antibody titers after CoronaVac priming and one-dose or two dose variant spike mRNA boosters. **f**, Neutralizing antibody titers after two-dose variant spike mRNA or protein boosters. Statistical significance was determined using the Wilcoxon rank sum test (a, c, d and f) or Wilcoxon signed-rank test (b and e). *p < 0.05, **p < 0.01, ***p<0.001, ****p<0.0001, and not significant (NS) p > 0.05.

**Extended Data Fig. 3.**
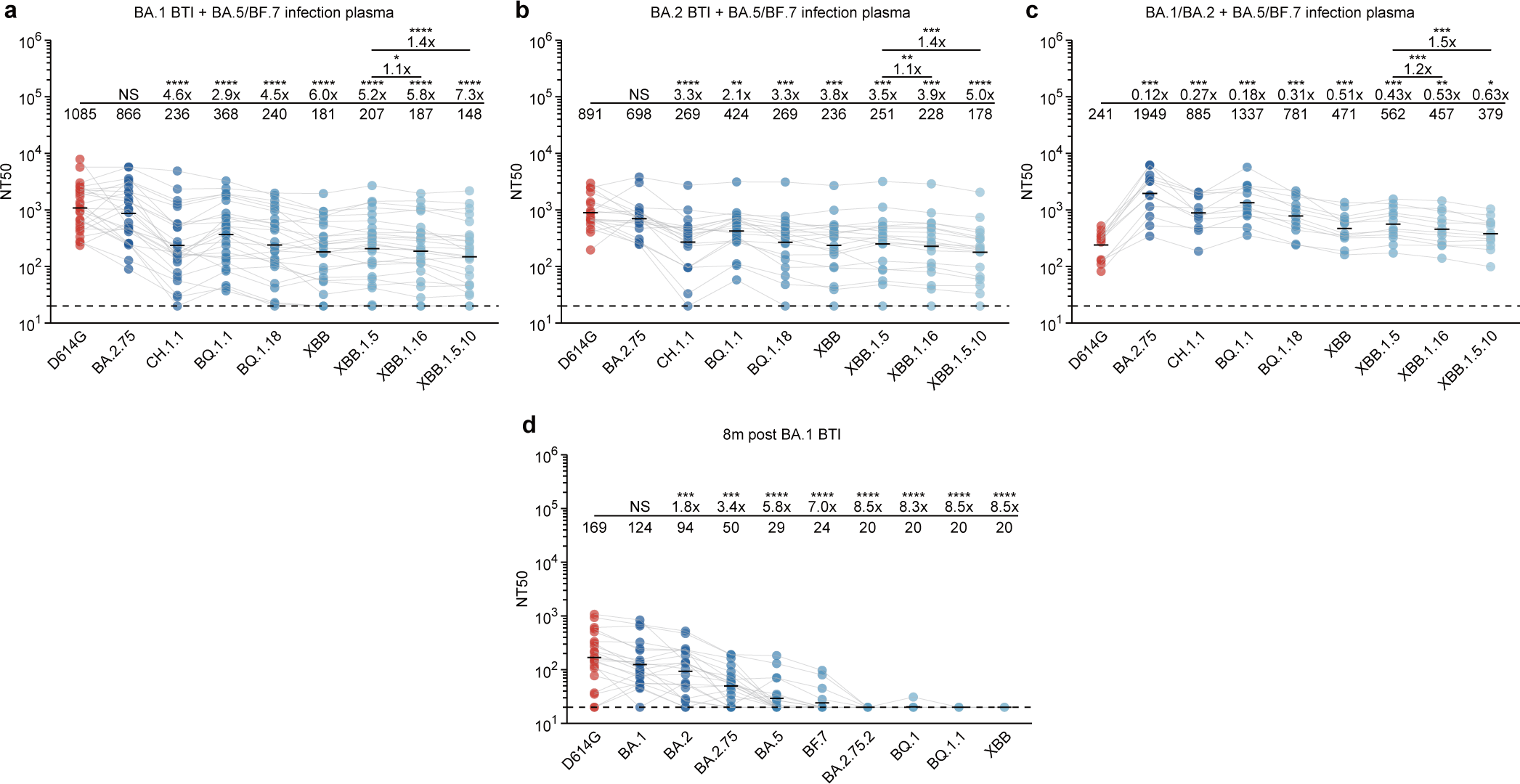
Antibody breadth of plasma after repeated Omicron infections. **a-d**, Plasma antibody titers against pseudotyped D614G and variants after (a) BA.1 BTI + BA.5/BF.7 infection (n = 26), (b) BA.2 BTI + BA.5/BF.7 infection (n = 19), (c) BA.1/BA.2 + BA.5/BF.7 infection (n = 12), d) 8 month post BA.1 BTI (n = 22). Fold changes between titers against variants and D614G were calculated and shown above the line. Statistical significance was determined using the Wilcoxon signed-rank test.

**Extended Data Fig. 4.**
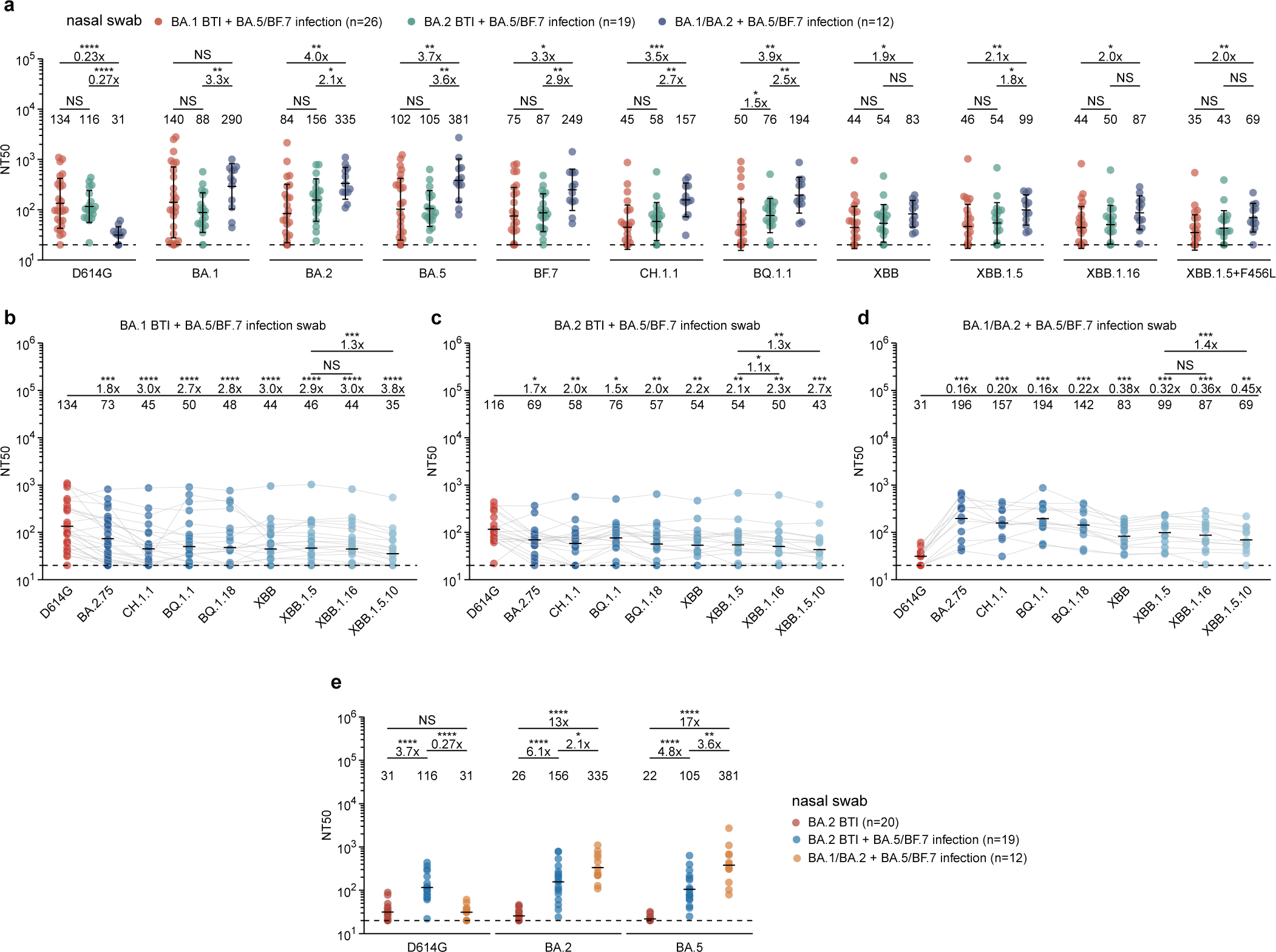
Neutralizing titers of nasal swabs after repeated Omicron infections. **a**, Comparison of nasal swab neutralizing titers among repeated Omicron infection cohorts. Nasal swab antibody titers against pseudotyped variants were measured. Fold changes between titers of different cohorts were calculated and shown above the line. Statistical significance was determined using the Wilcoxon rank sum test. **b**-**d**, Nasal swab antibody titers against pseudotyped D614G and variants after (**b**) BA.1 BTI + BA.5/BF.7 infection (n = 26), (**c**) BA.2 BTI + BA.5/BF.7 infection (n = 19), (**d**) BA.1/BA.2 + BA.5 infection (n = 12). Fold changes between titers against variants and D614G were calculated and shown above the line. Statistical significance was determined using the Wilcoxon signed-rank test in (b-d). **e**, Comparion of nasal swab antibody titers against pseudotyped D614G and variants among one-time breakthrough infection and repeated infection cohorts. Statistical significance was determined using the Wilcoxon rank sum test in (**e**). *p < 0.05, **p < 0.01, ***p<0.001, ****p<0.0001, and not significant (NS) p > 0.05.

**Extended Data Fig. 5.**
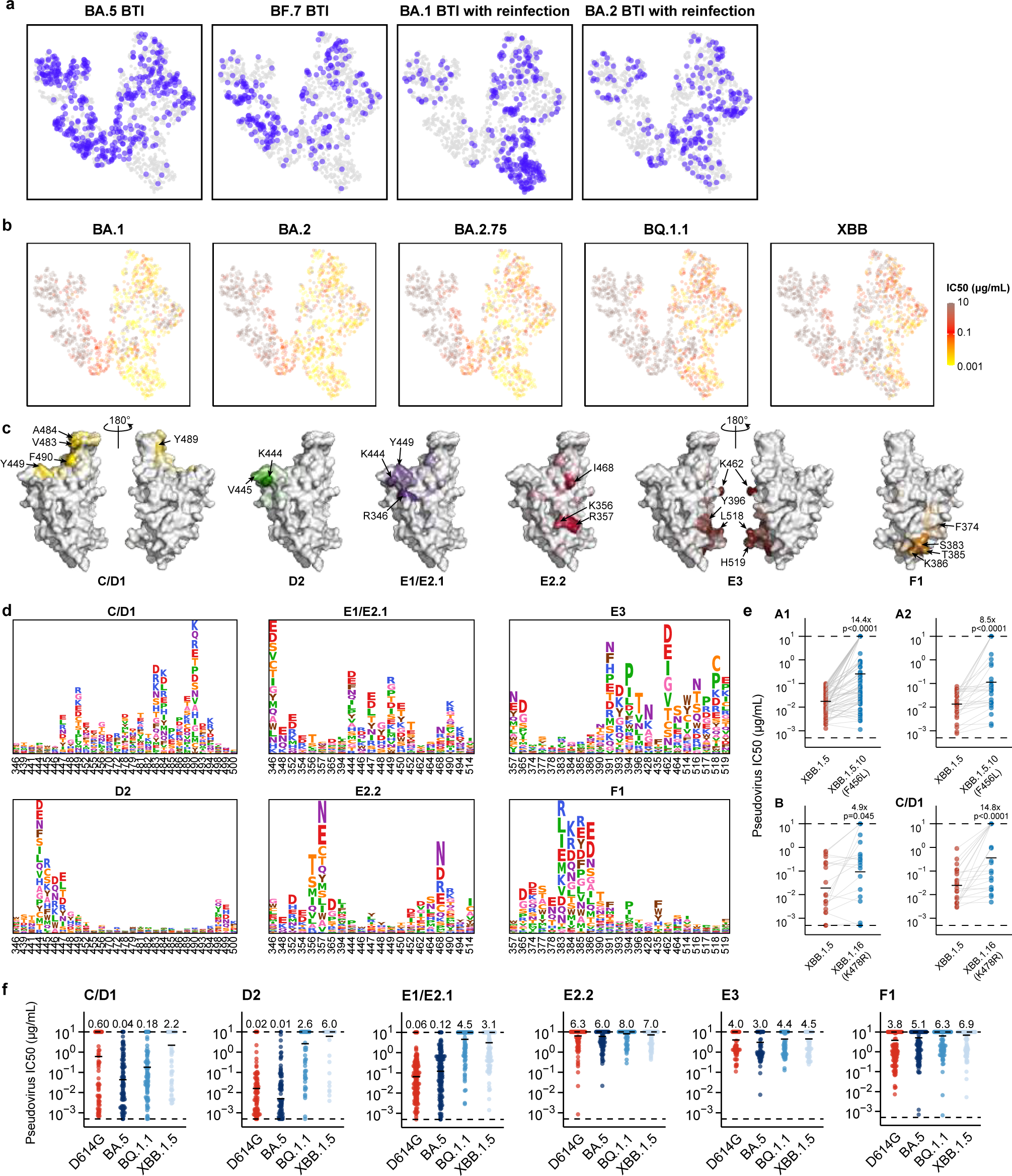
Characteristics of BA.5-reactive mAbs elicited by BA.5/BF.7 BTI or reinfection. **a,** Source of the antibodies are projected onto the UMAP embedding space. Antibodies from BA.5 BTI (n=445), BF.7 BTI (n=243), BA.1 BTI with reinfection (n=284), and BA.2 BTI with reinfection (n=232) are colored blue in the corresponding panel, and other antibodies are gray. **b**, Neutralization activities, denoted as IC50 values, against SARS-CoV-2 BA.1 (n = 1260), BA.2 (n = 1238), BA.2.75 (n=1238), BQ.1.1 (n = 1335) and XBB (n = 1341) spike-pseudotyped VSV are projected onto the UMAP embedding space. **c**, Average escape scores of epitope groups that are not shown in Fig. 4d (C/D1, D2, E1/E2.1, E2.2, E3, and F1) are illustrated on the structure model of the SARS-CoV-2 BA.5 RBD (PDB: 7XNS). Key residues with high escape scores for each group are labeled. **d**, Average DMS escape scores for these epitope groups are represented as sequence logos; residues are depicted using the standard one-letter code and colored based on their chemical properties. The height of each letter corresponds to the escape score of the respective mutation. **e**, Pseudovirus-neutralization activities of XBB.1.5-neutralizing mAbs in groups A1 (n=70) and A2 (n=23) against XBB.1.5 and XBB.1.5.10; and mAbs in groups B (n=15) and C/D1 (n=13) against XBB.1.5 and XBB.1.16. Fold changes in IC50 are labeled. P-values are calculated using two-tailed Wilcoxon signed-rank test of paired samples. **f,** Pseudovirus-neutralization activities of mAbs within the six crucial epitope groups (C/D1 [n = 76], D2 [n = 86], E1/E2.1 [n = 100], E2.2 [n = 124], E3 [n = 101], and F1 [n = 236]) are shown against SARS-CoV-2 D614G, BA.5, BQ.1.1, and XBB.1.5. Geometric mean IC50 values are displayed as bars and labeled above each group of data points.

**Extended Data Fig. 6.**
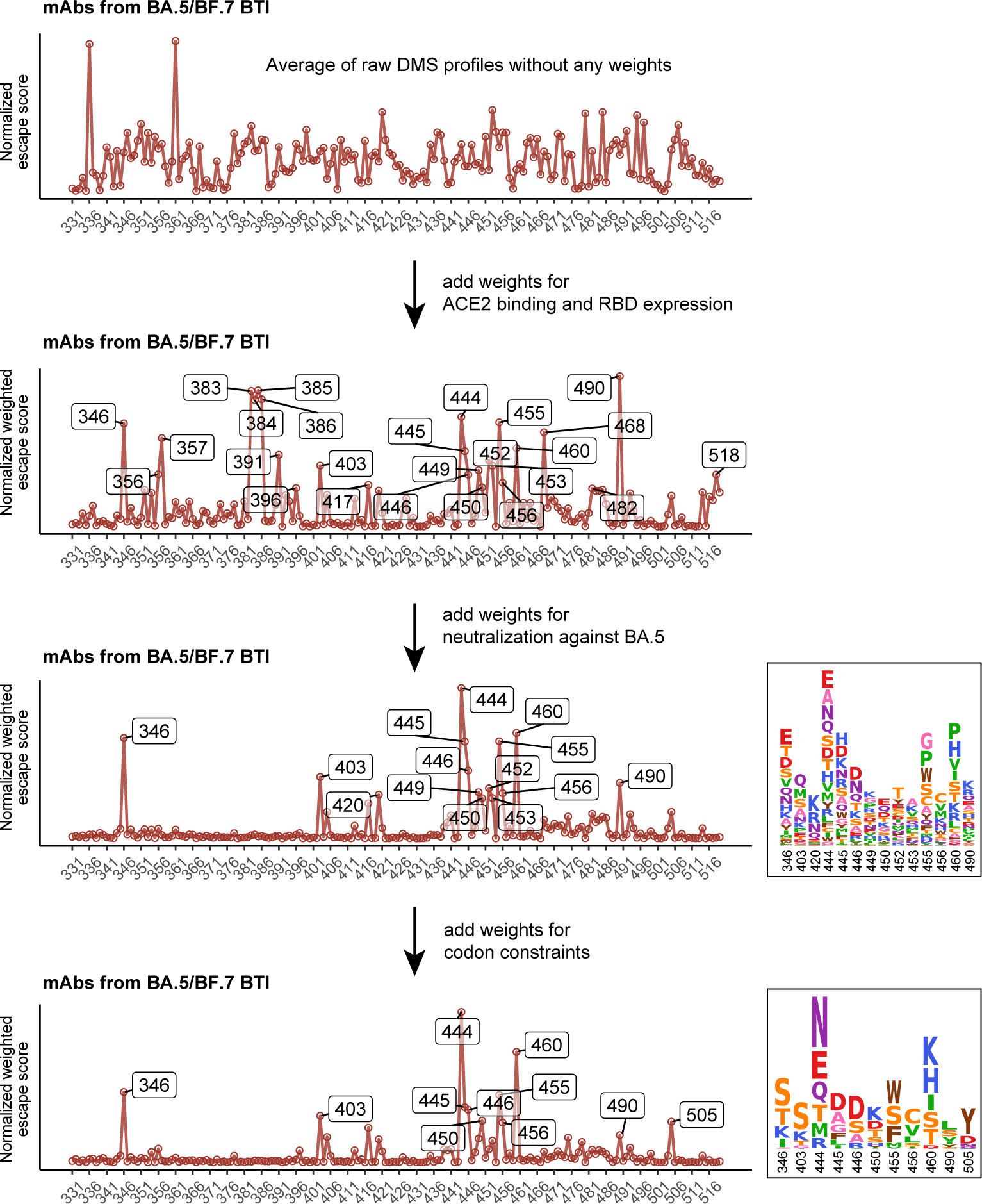
Workflow of calculating weighted escape scores of each mutation on RBD. Weights for ACE2 binding and RBD expression, neutralization activity, and codon usage are sequentially applied on the calculation to achieve informative results. Mutation preferences of BA.5 RBD under the pressure of NAbs from BA.5 or BF.7 BTI are shown.

**Extended Data Fig. 7.**
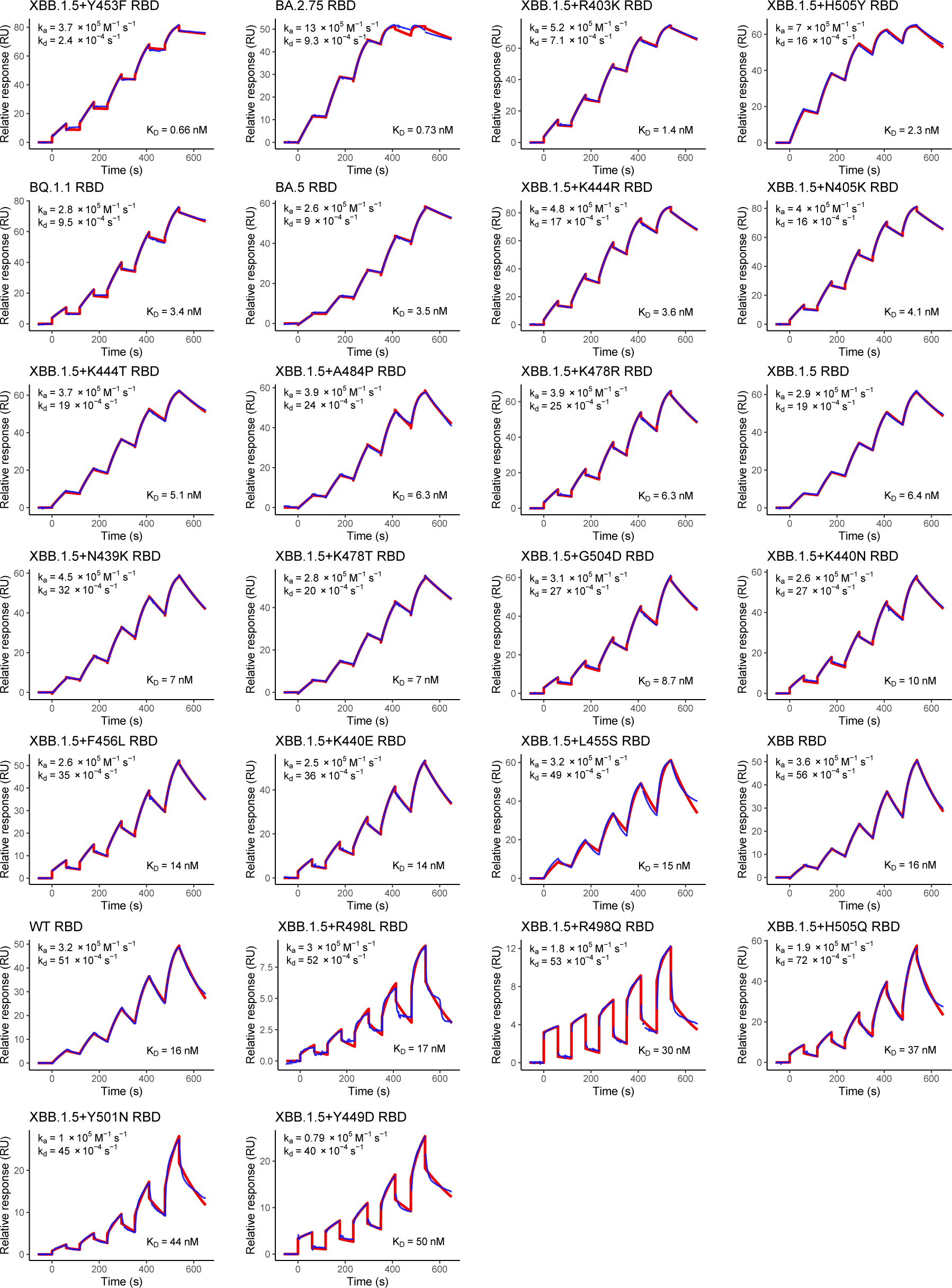
SPR sensorgrams for affinity of hACE2 and SARS-CoV-2 mutants RBD. Representative sensorgram of at least four replicates is shown for each RBD. Geometric mean kinetic constants k_a_, k_d_, and dissociation equilibrium constant K_D_ are labeled in each panel.

**Extended Data Fig. 8.**
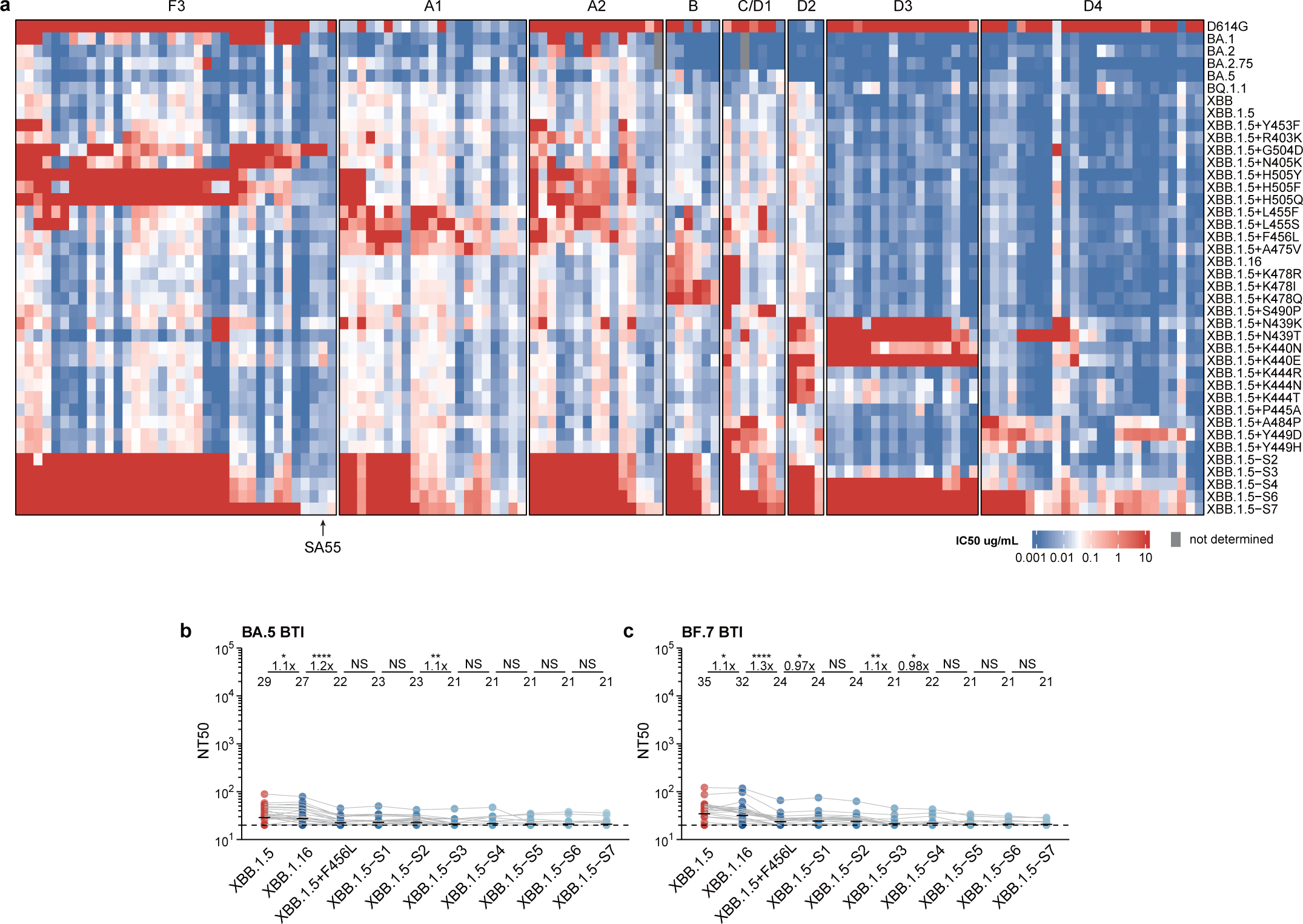
NAbs from BTI and reinfection are escaped by constructed mutants. **a**, IC50 values for representative potent XBB.1.5-neutralizing antibodies from different epitope groups against XBB.1.5 variants carrying individual or multiple escape mutations are shown. The order of antibodies is the same as that in Fig. 6c. **b**, Pseudovirus NT50 for SARS-CoV-2 XBB.1.5-based mutants are shown using plasma from convalescent individuals who experienced BA.5 (n=36) or BF.7 BTI (n = 30). Statistical tests are performed between neighboring mutants. P-values are calculated using two-tailed Wilcoxon signed-rank tests on paired samples. *p < 0.05, **p < 0.01, ****p < 0.0001, and p > 0.05 (NS).

